# Design principles to assemble drug combinations for effective tuberculosis therapy using interpretable pairwise drug response measurements

**DOI:** 10.1101/2021.12.05.471248

**Authors:** Jonah Larkins-Ford, Yonatan N. Degefu, Nhi Van, Artem Sokolov, Bree B. Aldridge

## Abstract

A challenge in designing treatment regimens for tuberculosis is the necessity to use three or more antibiotics in combination. The combination space is too large to be comprehensively assayed; therefore, only a small number of possible combinations are tested. We narrowed the prohibitively large search space of combination drug responses by breaking down high-order combinations into units of drug pairs. Using pairwise drug potency and drug interaction metrics from in vitro experiments across multiple growth environments, we trained machine learning models to predict outcomes associated with higher-order combinations in the BALB/c relapsing mouse model, an important preclinical model for drug development. We systematically predicted treatment outcomes of >500 combinations among twelve antibiotics. Our classifiers performed well on test data and predicted many novel combinations to be improved over bedaquiline + pretomanid + linezolid, an effective regimen for multidrug-resistant tuberculosis that also shortens treatment in BALB/c mice compared to the standard of care. To understand the design features of effective drug combinations, we reformulated classifiers as simple rulesets to reveal guiding principles of constructing combination therapies for both preclinical and clinical outcomes. One example ruleset is to include a drug pair that is synergistic in dormancy and another pair that is potent in a cholesterol-rich growth environment. These rulesets are predictive, intuitive, and practical, thus enabling rational construction of effective drug combinations based on in vitro pairwise drug synergies and potencies. As more preclinical and clinical drug combination data become available, we expect to improve predictions and combination design rules.

## Introduction

Tuberculosis (TB) remains a prevalent and important global health concern, with more than ten million people falling ill and about 1.4 million dying in 2019 (World Health Organization, 2020). Multiple drugs are used to treat TB because combination therapy shortens treatment duration, reduces disease relapse (increases treatment efficacy), and lowers the rate of drug resistance development compared with monotherapy (Fox et al., 1999). The standard of care (SOC) for TB treatment was developed almost 40 years ago and consists of four drugs (isoniazid (H), rifampicin (R), pyrazinamide (Z), ethambutol (E); HRZE) given for two months followed by two drugs (H and R; HR) given for another four to seven months (Fox et al., 1999). New multidrug therapies are needed to improve treatment outcomes and should include drugs that shorten treatment, increase treatment efficacy, or both.

Efforts to develop new antibiotics and combination therapies for TB have been highly productive (Aldridge et al., 2021). The large combination space created by the dozens of currently used drugs and the many under development (newtbdrugs.org) cannot be surveyed clinically. There is new evidence that improved drug combinations are in this space, as a recent clinical study identified a four-drug combination that shortened treatment by two months by substituting two drugs (H and R) from the SOC with moxifloxacin (M) and rifapentine (P) (Dorman et al., 2021). Furthermore, the combination consisting of bedaquiline (B), pretomanid (Pa) and linezolid (L), BPaL, has become an example for attainable TB treatment improvement in combination space because it shortened treatment of multidrug-resistant TB from over two years to six months with increased efficacy from less than 50% to more than 90% cure (Conradie et al., 2020). Reciprocal methods to clinical studies are needed to rapidly and systematically design combination therapies.

Preclinical animal studies are the primary tool in identifying drug combinations for clinical evaluation. The BALB/c relapsing mouse model (RMM) of durable treatment identified BPaL as a highly effective combination that showed faster and more effective curing than the three-drug mouse SOC (HRZ)(Xu et al., 2019). However, the number of combinations that can be tested in mouse studies is limited, and methods that prioritize drug combinations for preclinical testing are needed. We recently demonstrated that *in vitro* drug combination measurements in suites of multiple growth conditions were predictive of treatment improvement over the SOC in the RMM (Larkins-Ford et al., 2021). Those results suggest the potential to use *in vitro* measurements to select drug combinations with a high probability for treatment improvement over the SOC. However, the approach would still require measuring thousands of drug combinations to systematically survey the combination space, which is not feasible in either preclinical animal studies or *in vitro* experiments (Figure 1A).

**Figure 1.**
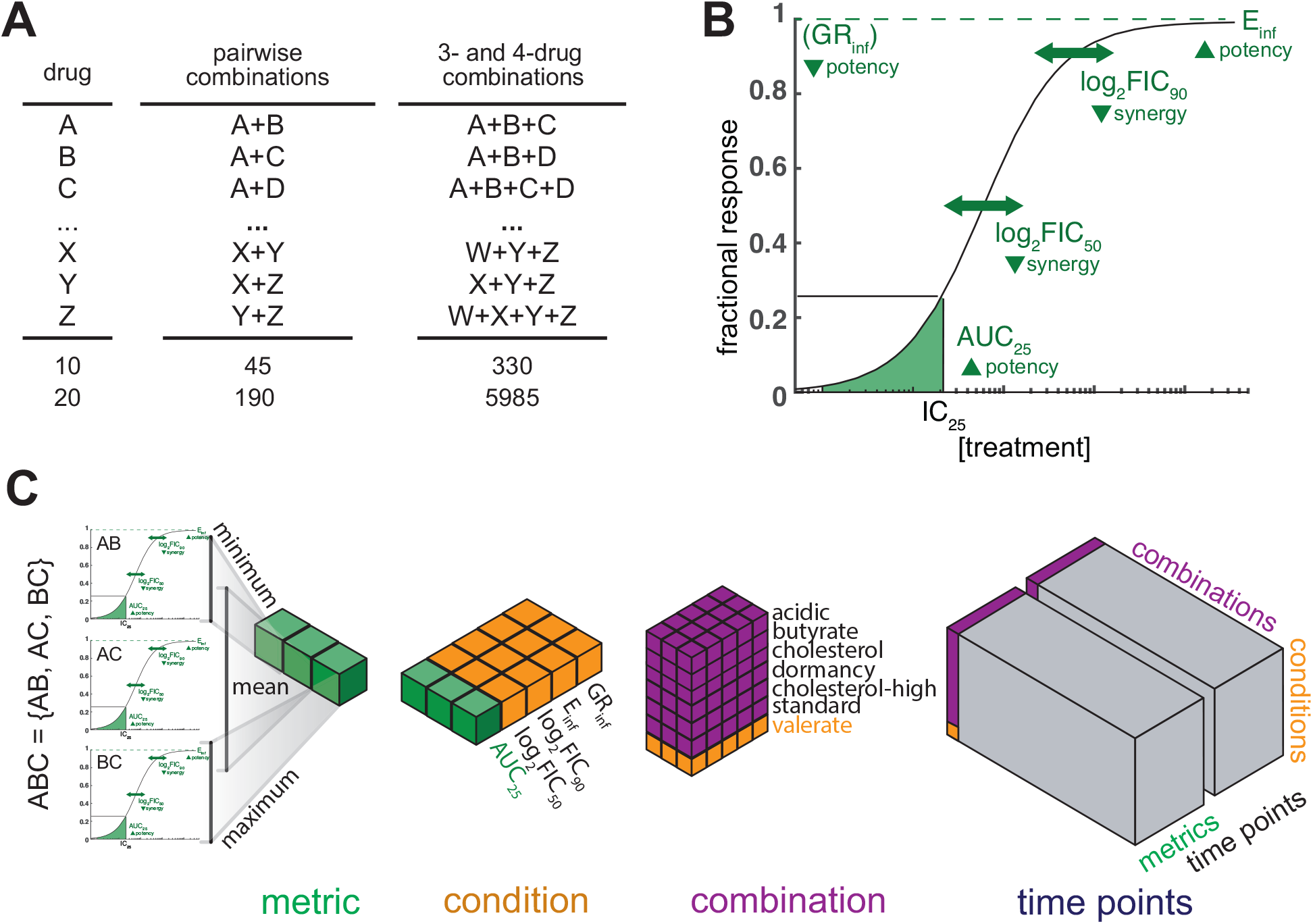
Data structure to organize *in vitro* drug pair data underlying higher-order drug combinations. (A) Summary of combinatorial explosion going from single drugs to three- and four-drug combinations for ten and twenty drugs. (B) Diagram of drug combination dose response curve, highlighting four (E_inf_, log_2_FIC_90_, log_2_FIC_50_, AUC_25_) of the five metrics calculated. GRinf is not diagramed because a separate dose response curve is used (Hafner et al., 2016). Below each metric is an arrow that points to whether low (down arrow) or high (up arrow) metric values are potent or synergistic. (C) Diagram of data structure used in the study. Combination ABC is composed of three drug pairs: AB, AC, BC. Metrics from each pairwise dose response curve are collated and summarized by calculating the minimum, maximum and mean for each metric (green) for every measured growth condition and time point. The summary metrics for a combination in an *in vitro* condition (orange) are compiled and concatenated with the metrics for all *in vitro* conditions (purple) to constitute all the pairwise data underlying a high-order combination at a given time point. The totality of data from all combinations (grey) at two time points in seven growth conditions and five metrics comprise the *in vitro* dataset used in this study.

One approach to searching the drug combination space more efficiently is to address the combinatorial explosion by utilizing drug pair data instead of requiring empirical data of three- and four-way combinations (Figure 1A). For example, there are almost 6,000 three- and four-drug combinations among twenty drugs but only 190 drug pairs; therefore, a method to understand and predict high-order combination treatment outcomes based on pairwise measurements alone would improve efficiency by ∼30-fold. The success of computational algorithms developed to predict the *in vitro* behavior of high-order drug combinations (three or more drugs) from the underlying low-order drug combinations (Wood et al., 2012, Katzir et al., 2019, Chandrasekaran et al., 2016, Cokol et al., 2017) indicates that pairwise drug interactions contain information important for understanding the activity of high-order drug combinations. These methods were developed to investigate drug interactions, which describe how drugs in combination can interact to produce combination effects that are greater than, less than, or as good as the effects of individual drugs (synergy, antagonism, and additivity, respectively). The prior success in mapping pairwise to high-order drug interactions *in vitro* suggests that it may also be possible to predict outcomes of multidrug therapies *in vivo* based on the properties of underlying drug pairs.

In this study, we develop a joint *in vitro* and computational framework to predict combination treatment outcomes for TB in mouse and clinical studies. Our study design is motivated by (a) the predictive signal of *in vitro* combination potencies and drug interactions to outcomes in the RMM using validated *in vitro* growth conditions, and (b) the ability to predict high-order drug interactions from underlying pairwise interactions. We used a dataset of systematic pairwise drug response data to develop machine learning models that accurately predict RMM and clinical outcomes of high-order combinations, creating a scalable and resource-sparing method to design combination therapies for TB rationally. We found that pairwise *in vitro* data carry a strong predictive signal, resulting in multiple rule sets for constructing effective high-order combinations based on the interaction and potencies of drug pairs as the building blocks. The resulting rule sets provide interpretable heuristics for rational assembly of optimized drug combinations that are better than the standard of care and BPaL in the RMM. We found that top combinations could be designed by assembling, for example, a pair that is synergistic in a culture environment that induces dormancy with another pair that is potent in a lipid-rich growth medium. Furthermore, the principles of combination design translated from the RMM to clinical outcomes. Our framework simultaneously creates accurate predictions of combination therapy outcomes in preclinical models and interpretable rules to construct optimized combinations.

## Results

### Organizing high-order drug combinations by summarizing pairwise drug combination data

We hypothesized that high-order drug combination RMM treatment outcomes could be predicted using *in vitro* pairwise drug combination measurements because pairwise drug interaction data are predictive of high-order drug interactions in *Mycobacterium tuberculosis* (Mtb), *Escherichia coli*, and cancer cells (Wood et al., 2012, Katzir et al., 2019, Chandrasekaran et al., 2016, Julkunen et al., 2020). In addition, we have previously shown that *in vitro* drug combination response data is predictive of RMM treatment outcomes (Larkins-Ford et al., 2021). To test this hypothesis, we designed a data structure to organize pairwise *in vitro* drug combination measurements across a range of drug pair potencies and drug interactions for each high-order drug combination under consideration (Figure 1).

We used pairwise drug combination response data from a large-scale study that contains *in vitro* measurement of two- and three-drug combinations among ten commonly used anti-tuberculosis drugs (Larkins-Ford et al., 2021). To broaden the scope of *in vivo* studies, we expanded this 10-drug set (bedaquiline, clofazimine, ethambutol, isoniazid, linezolid, moxifloxacin, pretomanid, pyrazinamide, rifapentine, rifampicin) with pairwise measurement to SQ109 and sutezolid for a total of 12 drugs. A portion of the SQ109 pairwise data was described previously (Egbelowo et al., 2021), while its remainder and all of the sutezolid data are new to this study. An equipotent mixture of each drug was measured at multiple doses to generate a pairwise dose response curve (Cokol et al., 2017, Larkins-Ford et al., 2021, Van et al., 2021). Drug combinations were measured in seven *in vitro* growth conditions relevant to the environments encountered by Mtb during infection: fatty acid carbon sources consisting of (1) butyrate, (2) valerate, (3) 0.05 mM cholesterol, and (4) 0.2 mM cholesterol (cholesterol-high), as well as (5) acidic medium (acidic), (6) non-replicating/hypoxic medium (dormancy), and (7) standard laboratory growth medium (standard). Longitudinal measurements were made, and two time points were targeted that represent a relatively consistent drug exposure time across conditions (constant), as well as the maximal drug exposure time relative to the doubling time of Mtb in each growth condition (terminal; constant and terminal times were the same for the standard condition). Five metrics were calculated for each dose-response curve (Figure 1B), capturing combination potency (AUC_25_, E_inf_, GR_inf_) and drug interactions at low and high dose levels (log_2_FIC_50_, log_2_FIC_90_). In total, 65 metrics were calculated for each of the 60 drug pairs, totaling 3900 pairwise dose response metrics (Table S1).

When breaking down high-order drug combinations into corresponding sets of drug pairs (e.g., ABC into AB, AC, and BC), some drug pairs will serve as components of multiple high-order drug combinations (e.g., AB is a component of ABC, ABD, ABCD). An important consequence is that each drug pair in a high-order drug combination will have an associated metric (e.g., for combination ABC, there will be an E_inf_ metric for AB, AC, and CD), but drug combinations of different orders will consist of different numbers of drug pairs and consequently have different numbers of pairwise dose-response metrics. To make combinations of different orders comparable, we devised a data structure where each high-order drug combination was represented by the same number of dose-response features, accomplished by aggregating the constituent pairwise metrics (AUC_25_, E_inf_, GR_inf_, log_2_FIC_50_, log_2_FIC_90_) using three summary statistics: minimum (min), maximum (max), and arithmetic mean (mean; Figure 1C). The three summary statistics ensured a uniform data structure of 195 features (mean, min, max of pairs for each metric, condition, and time point) from pairwise data for all high-order combinations (Table S2), facilitating downstream analyses.

### Pairwise data are predictive of high-order in vivo treatment outcome

To test the hypothesis that *in vivo* high-order drug combination treatment outcomes can be predicted from *in vitro* pairwise treatment data, we binned the dataset of high-order (three-, four-, and five-drug) combinations by assessing whether each combination was better (+C1) or worse (-C1) than the SOC in the RMM outcome using published animal studies. Combinations were deemed better if they achieved lower relapse (increased efficacy), similar relapse percentage with shorter treatment time (treatment shortening), or both, over the SOC (Table S3). Principal Component Analysis (PCA) revealed partial separation of +C1 and -C1 combinations along the first principal component (PC), indicating a strong predictive signal in pairwise data and suggesting that linear combinations of *in vitro* pairwise drug responses may be sufficient to distinguish drug combinations with different *in vivo* outcomes, even in the absence of trained supervised learning models. Notably, the signal was robust to the number of drugs involved in a combination, as we observed separation between 3-drug and 4+-drug combinations along the second PC, which was orthogonal to the first (Figure S1A).

Though PCA revealed partial separation of -C1 and +C1 combinations, the remaining overlap hinders accurate classification of candidate combinations using PCs alone, so we turned to supervised machine learning (ML) to increase classification accuracy. We evaluated seven ML algorithms for their ability to distinguish +C1 and -C1 combinations and compared their performance with repeated random partitioning of data for model training and evaluation. We observed ensemble methods, such as Random Forest (RF), to be top performers (Table S4), with corresponding classifiers achieving high (AUC > 0.86) accuracy on both training and test data. We, therefore, chose RF for all subsequent analyses.

By necessity, TB drug regimen development is iterative in that drugs are added to or substituted into effective combination scaffolds. Testing of combinations with new antibiotics often begins by adding or substituting a drug into a combination that has been previously tested. To simulate the process of using a new drug in combination with existing drugs, we treated each drug from the 12-drug dataset as a “new” drug. For this analysis, each drug was individually left out except for rifampicin, which could not be left out because too few -C1 combinations remained in the training set. We reserved combinations containing the candidate drug for testing (“leave-one-drug-out”) and trained a model on the remaining drug combinations. For each of the 11 “leave-one-drug-out” training/test sets, we included the HRZE combination (four-drug SOC) in the test set to evaluate combination prediction compared with the SOC. The models correctly predicted whether including the “left-out” drug improved treatment outcome (mean AUC +/- SEM 0.91 +/- 0.04, Table S5). Though the inclusion of any one drug into the scaffold was not a requirement for accurate performance, the exception was the model trained after bedaquiline was left out, which produced a random classifier (AUC = 0.58). Together, these results demonstrate that RMM outcomes of combinations containing a previously untested drug can be effectively predicted using pairwise *in vitro* measurements with minor drug-specific limitations.

### Additional classifiers predict top performing combination outcomes in vivo

Of all the possible 575 three- and four-drug combinations among the 12 drugs, only 39 (∼7%) were annotated with an RMM outcome, which we further split into 29 training and 10 test combinations. Given the SOC classifier (Figure 2A), we used the 29 annotated combinations in the training set to compute the optimal classification threshold (Youden’s J; P(+C1)=0.71) and applied it to categorize *in vitro* data from the 10 test combinations and the remaining 536 (∼93%) candidate combinations (Table S6). Of the 536 candidates, the classifier predicted 400 (76%) to be an improvement over the SOC, which was too high for effective follow-up. We, therefore, sought to prioritize the candidates further by asking whether they also outperform bedaquiline, pretomanid, and linezolid (BPaL). This combination is better than the SOC in the RMM (Xu et al., 2019, Mudde et al., 2021, Berg et al., 2021) and is a highly effective combination in the clinic, where it has been used to dramatically shorten the treatment time of multidrug-resistant TB (MDR TB) (Conradie et al., 2020).

**Figure 2.**
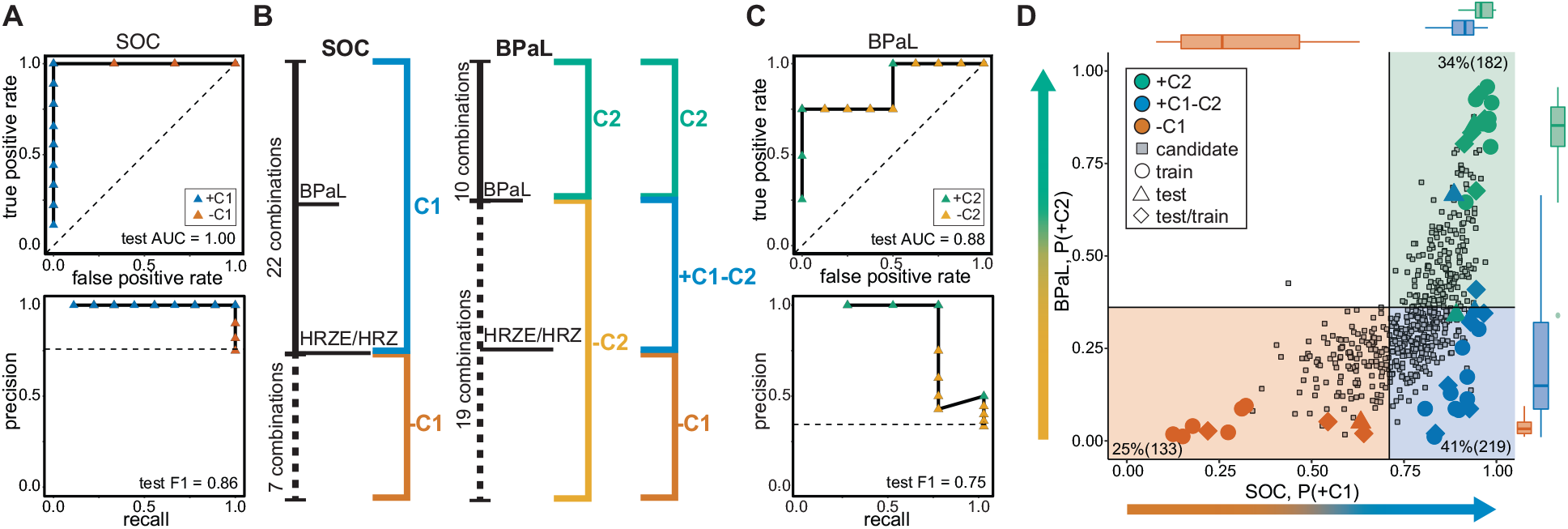
*in vitro* pairwise data is predictive of treatment improvement *in vivo*. (A) Receiver operator characteristic (ROC, upper panel) and precision recall (PR, lower panel) curves associated with an SOC random forest classifier trained using all summary pairwise features from seven *in vitro* growth conditions. The model was trained on 70% of annotated combinations and tested on the remaining 30%. Test combinations are colored by outcome annotation (blue = +C1, better than the standard of care; orange = -C1, standard of care or worse). (B) Schematic of combinations in the training set with annotations indicated by color and brackets. Selected combinations important for defining classes are indicated with single drug letter abbreviations (Table 1). (C) Receiver operator characteristic (ROC, upper panel) and precision recall (PR, lower panel) curves associated with a BPaL random forest classifier trained using all summary pairwise features from seven *in vitro* growth conditions. The model was trained on 70% of annotated combinations and tested on the remaining 30%. Test combinations are colored by outcome annotation (green = +C2, better than the BPaL; yellow = -C2, BPaL or worse). (D) Probability scatter plot for SOC model predictions (+C1 probability) and BPaL model predictions (+C2 probability). Marginal box plots show the annotated combination probability distributions. Annotated combinations are colored as in panel B (orange = -C1, blue = +C1-C2, green = +C2). Training combinations for SOC and BPaL are labeled with circles. Test combinations are labeled with triangles. Combinations that were used for training in SOC and testing in BPaL or testing in SOC modeling and training in BPaL modeling are labeled with diamonds. Combinations without annotations (candidates) are labeled with grey squares, and the number and proportion of candidates in three quadrants are indicated. A probability threshold associated with Youden’s J is shown with solid black lines. Probability regions showing classification of combinations within regions are colored as in panel B.

**Table 1.** Abbreviations used in this study. Abbreviations along with brief descriptions are listed.

| **Drugs** |  |
| --- | --- |
| **B** | bedaquiline, ATP synthesis inhibitor |
| **C** | clofazimine, antimycobacterial/multi-process inhibitor |
| **E** | ethambutol, cell wall synthesis inhibitor |
| **H** | isoniazid, cell wall synthesis inhibitor |
| **L** | linezolid, protein synthesis inhibitor |
| **M** | moxifloxacin, DNA synthesis inhibitor |
| **Pa** | pretomanid, cell wall synthesis inhibitor/ nitric oxide production |
| **Z** | pyrazinamide, antimycobacterial/multi-process inhibitor |
| **R** | rifampicin, transcriptional inhibitor |
| **P** | rifapentine, transcriptional inhibitor |
| **Su** | sutezolid, protein synthesis inhibitor |
| **Sq** | SQ109, multi-process inhibitor |
| **Drug Combination** |  |
| **BPaL** | bedaquiline + pretomanid + linezolid |
| **HRZE** | isoniazid + rifampicin + pyrazinamide + ethambutol - four-drug<br>standard of care |
| **BPaMZ** | bedaquiline + pretomanid + moxifloxacin + pyrazinamide |
| **HRZM** | isoniazid + rifampicin + pyrazinamide + moxifloxacin |
| **MRZE** | moxifloxacin + rifampicin + pyrazinamide + ethambutol |
| **HPZM** | isoniazid + rifapentine + pyrazinamide + moxifloxacin |
| **HRZ** | isoniazid + rifampicin + pyrazinamide |
| **Treatment outcome and classification** |  |
| **+C1** | better than standard of care |
| **-C1** | as good or worse than the standard of care (HRZE or HRZ) |
| **+C2** | better than BPaL |
| **-C2** | as good or worse than BPaL |
| **+C1-C2 (+C1 and -C2)** | Overlap between better than standard of care and worse than BPaL |
| **SOC** | Standard of Care |
| **TTP** | Time to Culture Positivity |
| **Mouse model** |  |
| **RMM** | Relapsing mouse model |
| **In vitro models** |  |
| **a** | acidic |
| **b** | butyrate |
| **c** | cholesterol (0.05 mM) |
| **d** | dormancy |
| **h** | cholesterol-high (0.2 mM) |
| **s** | standard |
| **v** | valerate |
| **Data and Metrics** |  |
| **C** | constant time point |
| **T** | terminal time point |
| **CT** | constant and terminal time points are the same |
| **Log<sub>2</sub>FIC<sub>n</sub>** | fractional inhibitory concentration at n % growth inhibition |
| **AUC<sub>25</sub>** | the normalized area under the dose-response curve to the 25% inhibition point |
| **E<sub>inf</sub>** | the effect at infinite drug concentration (maximum achievable effect) |
| **GR<sub>inf</sub>** | normalized growth inhibition effect at infinite drug concentration (maximum achievable effect) |
| **ROC** | receiver operator characteristic |
| **AUC** | the area under the ROC curve |
| **PR** | precision-recall |
| **F1** | the harmonic mean of the precision and recall |
| **Machine Learning acronyms** |  |
| **PC** | Principal Component |
| **PCA** | Principal Component Analysis |
| **RF** | Random Forest |
| **DT** | Decision Tree |

We reannotated the RMM outcome (Figure 2B) according to whether it was better than BPaL (+C2) or not (-C2, Table S3). The +C2 group is a proper subset of the SOC +C1, and the new - C2 class combines the remaining +C1 (now labeled +C1-C2) and the previously labeled -C1 combinations. As with SOC, *in vitro* pairwise data are separated by the +C2 and -C2 labels along the top principal component (Figure S1B).

Using the same validation process we performed with SOC classifiers, we evaluated the performance of a model trained with features from all conditions for its ability to distinguish +C2/-C2 combinations. We observed comparable performance for the all-condition model during model training (AUC = 0.83) and high performance using the held-out test set (AUC = 0.89, Figure 2C). We also observed that combinations predicted to be better than BPaL (Youden’s J; P(+C2)>0.36) also tend to have the highest likelihood to improve treatment outcome over the SOC (P(+C1)>0.78, Figure 2D, Table S6). However, the converse is not true: high probability +C1 (P(+C1)>0.71) combinations may or may not be better than BPaL. This suggests that the SOC and BPaL classifiers are non-redundant, and classification for improvement over BPaL (182 combinations, 34%, Table S6) can further refine the set of +C1 combinations for experimental follow-up. We observed a wide range of probabilities for the BPaL classification (P(+C2) between ∼0.35 and ∼0.75) in which there are few annotated combinations; therefore, further prioritization may be achieved using a more conservative BPaL classification threshold (e.g., P(+C2)=0.5, 14% (73) +C2 combinations) or by ranking candidate combinations using probabilities (Figure 2D). Whichever method is used for candidate prioritization, we predict that there are many potential treatment-improving combinations using existing anti-tuberculosis drugs and that +C2 combinations represent a unique subset of treatment improving combinations.

### RMM outcome prediction is improved using subsets of in vitro conditions

Mtb encounters many environments during infection, and some are thought to contribute more than others to the requirement for long treatments. Models constructed from multiple conditions as a “sum-of-parts” are likely to be the most predictive because they represent the diversity of microenvironments encountered during an infection (Larkins-Ford et al., 2021). We asked which of the seven *in vitro* models were most predictive and whether a smaller set of *in vitro* conditions could be used to model RMM outcomes. We reasoned that a model trained with three conditions would be sufficient to represent the diversity of physiological states during infection while also constituting a practical set of experiments to perform. We also confirmed that increasing the number of conditions in a model beyond three did not generally improve performance (Figure S2).

After evaluating all possible three-condition models, we observed that all but one model was high performing for both outcomes (AUC > 0.7, Figure 3A) and that many performed better than the seven-condition model (SOC AUC=0.83, BPaL AUC=0.83). These results demonstrate that three conditions were sufficient to train models that were as good or better than a model trained on all possible condition information. We confirmed the high performance of three-condition models using test data and predicted candidate combination classification comparable to the all-condition model (Figure S3). Furthermore, high-performing models can be trained using many aggregated sets of three conditions (Figure S2). Finally, the high performance of three-condition sets for both BPaL and SOC outcome models suggests that using one of the two is sufficient for classifier evaluation. Therefore, we focused on only BPaL outcome models in subsequent analyses.

**Figure 3.**
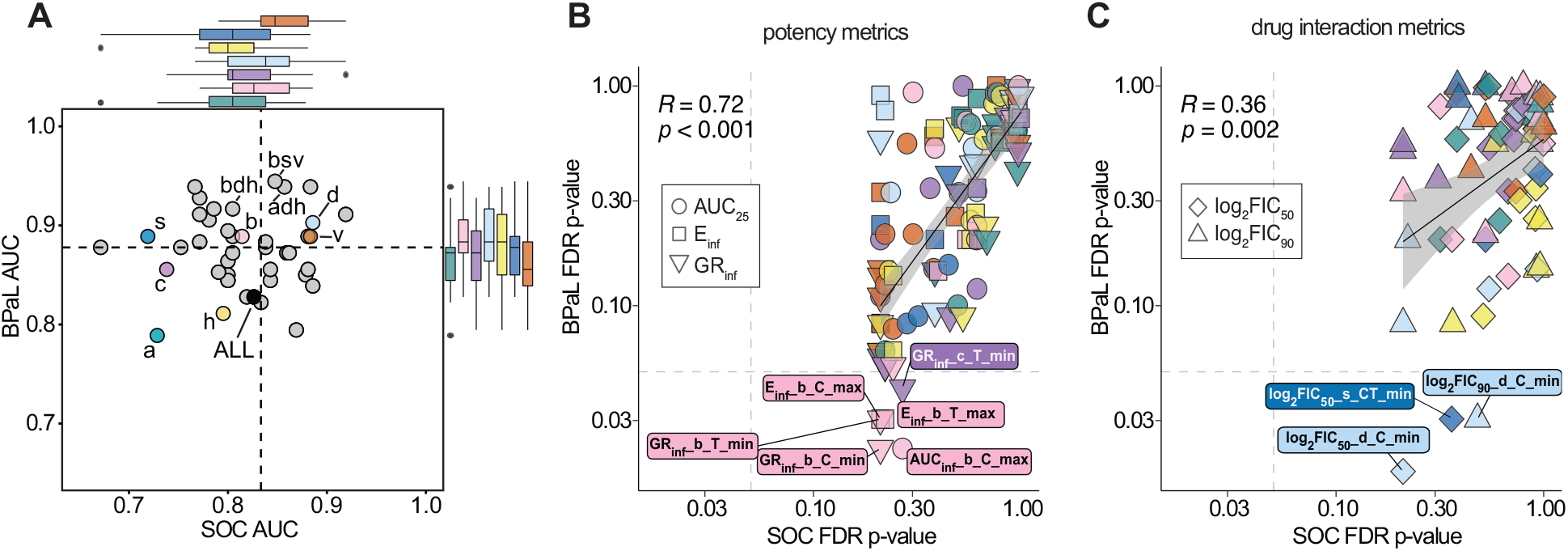
Predictive information in subsets of *in vitro* conditions and dose response metrics. (A) Scatter plot of model performance (AUC) for the SOC and BPaL machine learning models trained on data from one, three or all (seven) conditions and evaluated in cross-validation (see Figure S2 for performance of models from any number of conditions). Marginal boxplots indicate the performance of models containing each condition. Dashed line indicates the median performance across all models. One-, seven-, and selected three-condition subsets are labeled. Single condition abbreviations: a, acidic; b, butyrate; c, cholesterol; d, dormancy; h, cholesterol(high); s, standard; v, valerate (Table 1). (B) Scatter plot of p-values from the Wilcoxon rank-sum tests contrasting values of individual potency features across SOC (-C1 vs. +C1) and BPaL (-C2 vs. +C2) outcomes. Features are colored by *in vitro* condition and shaped by metric type (circle, AUC_25_; square, E_inf_; downward triangle, GR_inf_). P-values are corrected for multiple hypothesis testing within each outcome group (e.g., corrected for SOC comparison separate from BPaL comparison). Dashed lines show p=0.05. Features with FDR p-values <0.05 are annotated with extra information such as time (C or T for constant or terminal, respectively) and the summary statistic type (minimum, mean, or maximum). Linear regression line (solid black), confidence interval (shaded region), Pearson correlation coefficient (R) and associated p-value are indicated on plot. (C) Scatter plot of p-values from the Wilcoxon rank-sum tests contrasting values of individual drug interaction features across SOC (-C1 vs. +C1) and BPaL (- C2 vs. +C2) outcomes. Plot elements are analogous to those in panel B. Features are shaped by metric type (upward triangle, log_2_FIC_90_; diamond, log_2_FIC_50_).

Conditions that have important information for predicting *in vivo* outcomes should be those that improve model performance when included, even if the condition is not the highest performing when considered by itself. Therefore, we compared model training performance with and without each of the seven conditions. We expected that if a condition is sufficiently informative, a majority (>50%) of the models that include it should have increased performance compared to when that condition is excluded. We observed that 65% of models saw an increase in AUC when butyrate was included, with similar trends for dormancy (54% of models) and cholesterol-high (51% of models, Figure S4A). The trend towards increased performance was maintained among models using data from four or more conditions that included butyrate+dormancy+cholesterol-high compared to those with only two or fewer of these conditions (p=0.206, Figure S4B). Lastly, we observed that the model butyrate+dormancy+cholesterol-high was the sixth-highest performing three-condition model for the BPaL outcome (AUC=0.92, F1=0.81, Figure 3A, Table S7), and the top five three-condition models included at least one of these three conditions.

Though dormancy and butyrate were the top two highest-performing single-condition models (Table S7), the cholesterol-high condition performed modestly as a single-condition model compared to other growth environments. Nevertheless, models with other conditions improved upon the addition of cholesterol-high measurements (Figure S4A), suggesting that the condition carries an orthogonal signal to other conditions.

This analysis demonstrates that there is predictive information in many of the *in vitro* models, with some conditions carrying redundant information, while others provided an orthogonal signal that improved classifier performance. Future work to prioritize combination therapies based on pairwise measurement will therefore not require exhaustive measurement in many growth conditions but can instead focus on sets of three *in vitro* models with established predictive accuracy.

### Treatment outcome is driven by exceptional drug pairs rather than averaged pairwise properties

Evaluation of growth condition contribution to classifier performance provides one angle of interpretability, whereas further insight may be gained from evaluating individual features. We sought to understand what features could accurately distinguish +C2/-C2 and +C1/-C1 combinations and what feature values constituted +C2 combinations. We examined the values of individual features among all conditions and found that several correlated with the +C2/-C2 outcome class (9 of the 186 (∼5%) features; p<0.05, Wilcoxon rank-sum test, using Benjamini-Hochberg multiple hypothesis correction; Figure 3B and C, Table S8, Figure S5A). Though no features differed significantly between +C1 and -C1 drug combinations (all p>0.05), we nevertheless observed a strong correlation between the significance of potency features in the SOC and BPaL comparisons (Figure 3B, Pearson correlation, R=0.72, p<0.001). In contrast, there was a substantially weaker correlation in the importance of drug interaction metrics between the SOC and BPaL outcome thresholds (Figure 3C, Pearson correlation, R=0.36, p=0.002). As with potency features, several drug interactions features differed significantly between +C2 and -C2 combinations, but not between +C1 and –C1 ones (Figure 3C); this is consistent with a prior study where we found that RMM outcomes relative to the SOC were predicted by potency metrics rather than synergies (Larkins-Ford et al., 2021).

We noted that many significant features were from the butyrate and dormancy conditions (Figure 3B and C), supporting the use of a three-condition model that includes these conditions. We also observed that all of the significant features to correlate with the +C2/-C2 dichotomy describe the most potent and most synergistic pairs (e.g., minimum GR_inf_ and log2FIC values and maximum E_inf_ and AUC_25_ values among the underlying pairs of a high-order combination). These results suggest that a small number of strong drug pairs contribute more information about treatment improvement of a high-order combination than the average behavior of all involved pairs.

Furthermore, these observations are not specific to the training set and generalize when test combinations were also considered (Figure S5B and C). Taken together, these results indicate that the degree of treatment improvement of a drug combination (over BPaL and SOC) can be predicted using *in vitro* measurements of pairwise drug potency and that there are drug pair synergies when Mtb are dormant, which distinguish drug combinations that are better than BPaL.

### Design principles for constructing effective drug combinations

Given our observations that highly effective drug pairs are driving the treatment outcome of high-order drug combinations (Figure 3), we aimed to understand how to identify and compile effective drug pairs using *in vitro* measurements. Our goal was to compose a set of rules to guide the rational design of high-order drug combinations using drug pairs as the building blocks.

Decision tree classification mirrors human decision-making and can define a set of rules for classification tasks. To make the rules defined by the decision tree interpretable and straightforward, we focused on the features from the butyrate, dormancy, and cholesterol-high conditions, motivated by the largest increase in performance when these three conditions were included in a machine learning model. We selected a single potency and drug interaction feature from each condition by choosing features with the strongest association (lowest p-values) with the +C2/-C2 dichotomy based on the Wilcoxon rank-sum test analysis (Table S8). We used the training and test data split from the BPaL RF classifier and trained a decision tree (DT1) to identify the features and thresholds that were most informative for identifying +C2 combinations (Figure 4A). The rules defined by these features indicate that the first step in constructing a combination is to choose a potent drug pair in butyrate (GR_inf_ in butyrate at the constant time point < -0.38) and then choose a pair that is additive/synergistic in cholesterol-high (log_2_FIC_50_ in cholesterol-high at the terminal time point < 0.13). Several candidate drug combinations were also identified using these rules as likely to be +C2. The lower complexity of a two-feature decision tree yields did not alter accuracy when predicting the test set outcome compared to the RF classifier (83%), demonstrating that the simplicity of a short rule set provides an accurate understanding of how to construct effective combinations based on minimal information from pairwise measurement *in vitro*.

**Figure 4.**
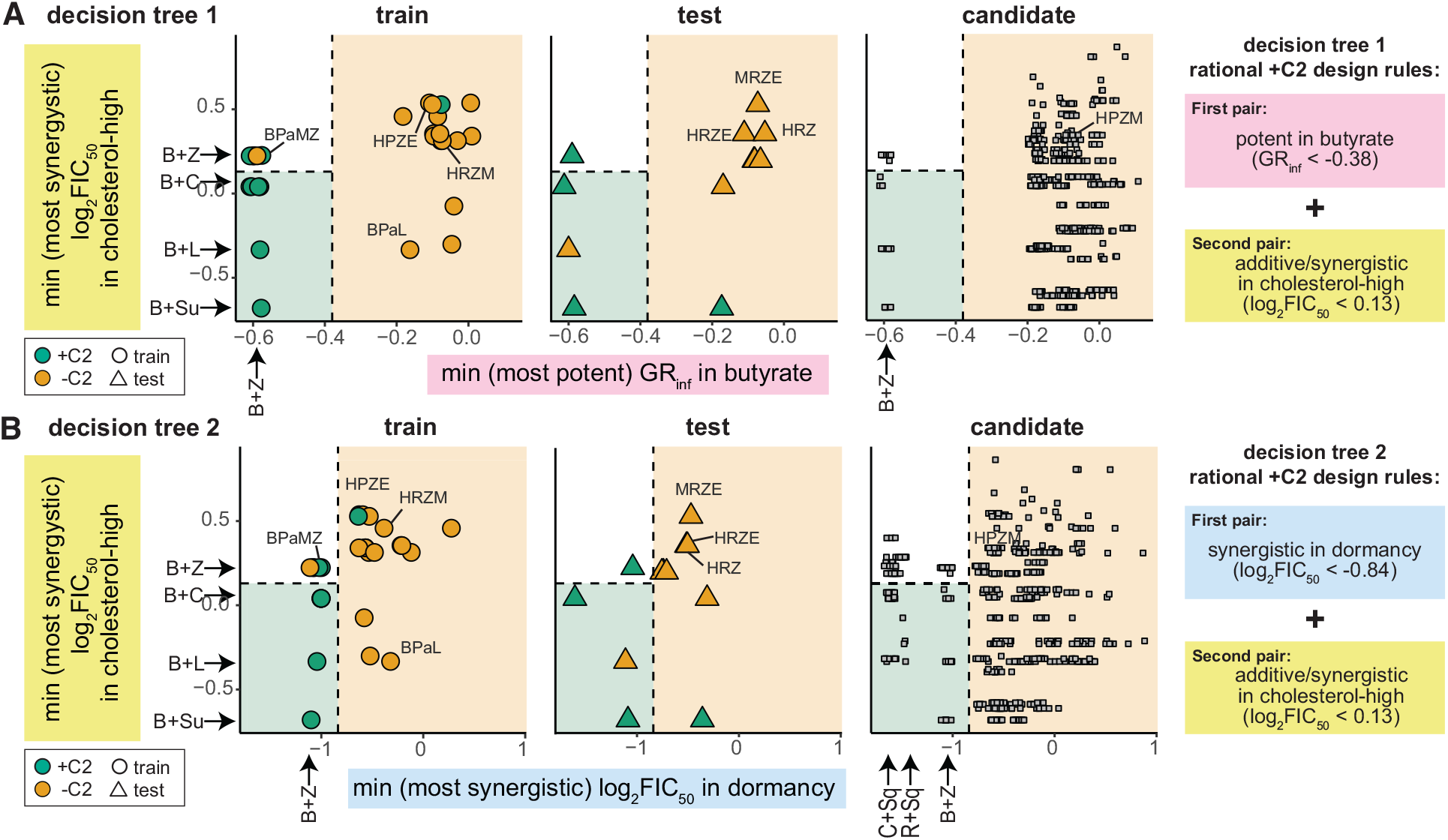
Rule sets for assembling +C2 (RMM) drug combinations based on effective drug pairs. Scatter plots of two metrics identified to be important for outperforming BPaL, shown as decision tree 1 (A) and an alternative decision tree 2 (B). Combinations are colored by classification (green = +C2, orange = -C2). Combinations are plotted separately based on whether they were used in decision tree model training (circle, left), testing (triangle, middle), or are candidates (square, right). Selected drug combinations are indicated with labels. Regions of the plot are colored based on the decision tree classification using thresholds (dashed lines) learned during training. White region denotes satisfying rule one but not rule two criteria for +C2 classification. Metric values of selected drug pairs are indicated along plot margins. Rules are written in logic format on the right

The first decision tree (DT1) used only two features of the many that were observed to separate +C2 from -C2 combinations, suggesting that we may be able to write other rule sets. We trained a second decision tree (DT2), with an emphasis on dormancy and cholesterol-high features. DT2 was observed to be similar to DT1, with the second rule (of additivity/synergy in cholesterol-high) being identical in both trees. Conversely, the first decision in DT2 is based on having a highly synergistic drug pair in dormancy (Figure 4B, log_2_FIC_50_ at the constant time point < - 0.84) instead of a potent pair in butyrate (from DT1). The ability to substitute the first rule with another shows redundancies in predictive signals among the metrics in the *in vitro* dataset. Taken together, the DTs define a set of interpretable rules that can govern the rational design of effective high-order drug combinations. Notably, the rule sets are not absolute. Multiple rule variations can instruct the design of effective +C2 therapies, guided by the availability of the conditions used for the pairwise *in vitro* measurement (for more DTs in other condition subsets, see Figure S6).

We used DT1 and DT2 to predict classification as +C2 or -C2 on candidate combinations (Figure 4). Many drugs and drug combinations were over-represented in the combinations predicted to be +C2 (47 drugs and combinations by Fisher’s exact test, p<0.05, after Benjamini-Hochberg multiple hypothesis correction, Table S9). Notably, we observed enrichment of combinations that include bedaquiline (B), pyrazinamide (Z), clofazimine (C), and SQ109 (Sq), suggesting that these drugs partner well with other drugs. Prominent in these over-represented combinations is bedaquiline + pyrazinamide (B+Z); this may be explained by how well B+Z satisfies one rule in each DT (potent in butyrate and synergistic in dormancy). However, the likelihood of high-order combinations that include B+Z to be +C2 increased when another additive or synergistic pair in cholesterol-high is also included in the combination (Figure 4A). Stated another way, if B+Z satisfied the first rule (potent in butyrate or synergistic in dormancy), a combination would be +C2 (green region) if a different pair contributed to the second rule (non-antagonism in cholesterol-high). We trained alternative DTs for other top 3-condition models (Figure S6). We observed that potent pairs in dormancy, butyrate, and standard medium and synergistic pairs in dormancy, cholesterol-high, and valerate are features of +C2 combinations. We also observed that a rule set might include both synergy (a synergistic pair in dormancy) and antagonism (mean behavior of antagonism among the pairs in acidic medium) (Figure S6D); therefore, synergy as a heuristic may be specific to the growth condition and whether a dominant drug pair or average pairwise drug interaction are considered.

We conclude that when *in vitro* pairwise data are predictive of combination treatment outcomes *in vivo*, simplified and intuitive heuristics can be developed to define and interpret design principles on how to construct combinations from the bottom up. A rules-based approach will enable us to glance at systematic pairwise drug response metrics in Mtb to optimize combination therapies without running classifiers. To fully realize the potential of our drug combination dataset and aid in this “at-a-glance” approach to combination building, we have provided heatmaps of key pairwise drug combination metrics (Figure S7).

### Translation of combination drug design principles to clinical outcomes

The effectiveness and interpretability of the classifiers and rules for rationally designing combinations support the utility of the presented *in vitro* dataset for understanding the drivers of drug combination efficacy in preclinical mouse models of durable treatment outcomes. In principle, this methodology is agnostic to the *in vivo* outcomes that the models will be trained on, so long as the *in vitro* conditions are predictive of the pharmacodynamics at the sites of infection. We next asked whether our *in vitro* data could inform our understanding of clinical outcomes of drug combination treatment. We compiled a list and scored the outcome of drug combinations that had been evaluated for bactericidal activity in phase 2a and phase 2b clinical trials (Table S3). Clinical outcomes were scored relative to the SOC, as BPaL is not yet known to be treatment shortening relative to the SOC for drug-sensitive Mtb in clinical studies (in contrast to the RMM). Consistent with previous studies (Rosenthal et al., 2012, Dorman et al., 2012, Bartelink et al., 2017, Gillespie et al., 2014, Li et al., 2015, Lanoix et al., 2016a) we observed some discordance in the classification of the effectiveness of drug combinations between RMM and clinical outcomes (Figure 5A, Table S3). This discordance is expected because the outcomes (bactericidal vs. relapse) are different and suggest that a model trained for SOC (RMM) may not necessarily predict combinations with bactericidal efficacy in clinical studies. Two of the six discordant combinations were HRZM and MRZE; both failed to improve HRZE in the ReMOX clinical trial (Gillespie et al., 2014) and are -C1 in our clinical annotation. We previously annotated both combinations for bactericidal activity in the BALB/c mouse model as -C1 (Larkins-Ford et al., 2021), suggesting that the source of discordance may be the difference in outcome type. Due to the high cost of misidentifying combinations for follow-up in clinical trials, developing models and rules that identify potentially treatment improving combinations in clinical trials, separate from the preclinical predictions, is highly important.

**Figure 5.**
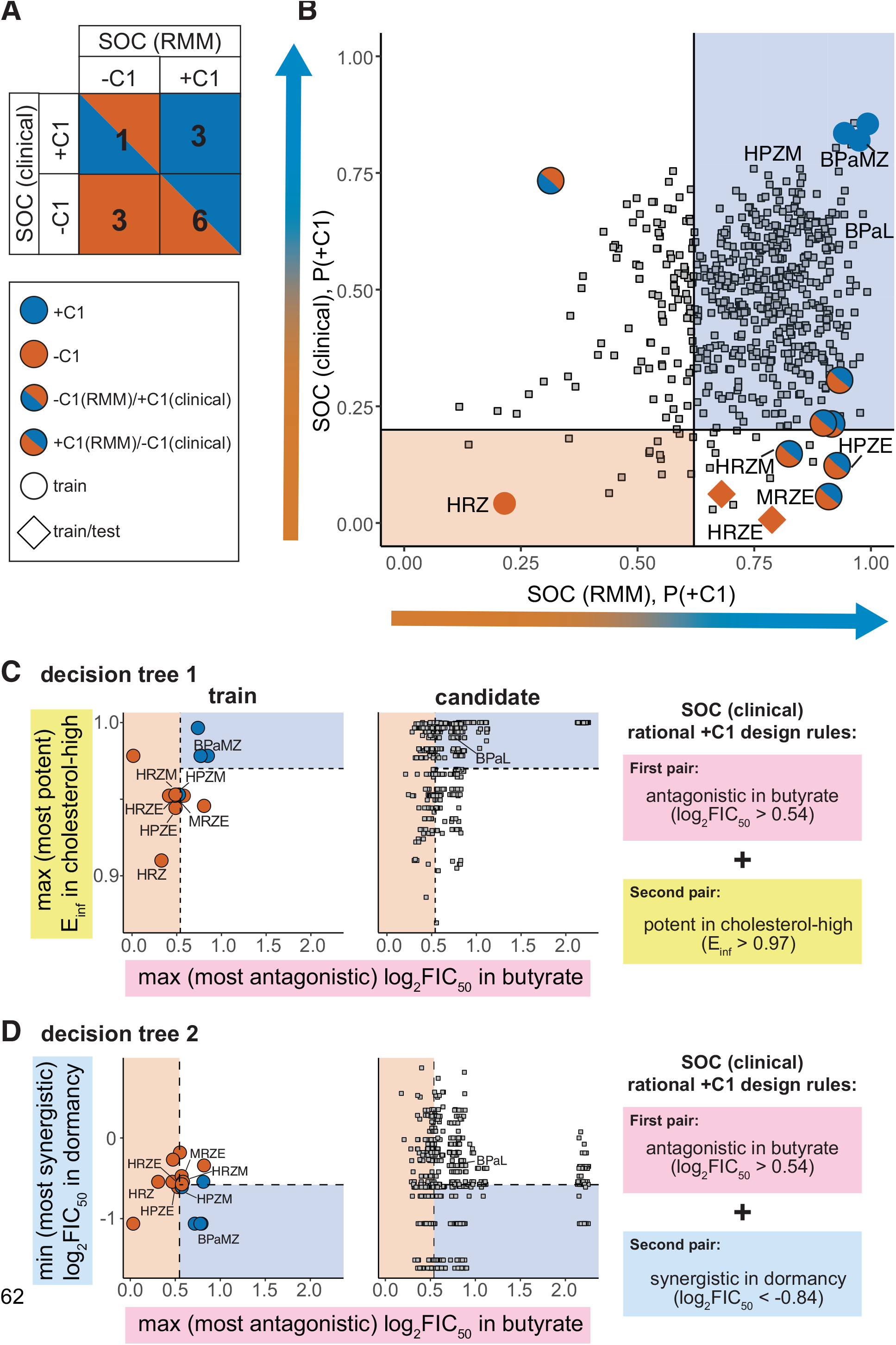
Modeling and rational design principals applied to clinical SOC outcome. (A) Overlap in +C1(blue) and -C1(red) drug combination categorization between SOC (RMM) and SOC (clinical) outcomes. Blue/red squares highlight differences between outcome annotation. (B) Probability scatter plot for SOC model predictions (+C1 probability) and clinical model predictions (+C1 probability) using the butyrate+dormancy+cholesterol-high condition data. Annotated combinations are colored as in panel A. Model training combinations for both SOC and clinical are labeled with circles. Combinations used for testing the SOC model and training the clinical model training are labeled with diamonds. Candidate combinations (without annotations) are labeled with grey squares, and the number and percent of candidates in three quadrants are indicated. Probability regions and threshold are labeled as in Figure 2 with regions colored as in panel A. (C) and (D) show scatter plots and alternative rule sets for two metrics identified as important for outperforming the SOC for decision tree 1 (C) and decision tree 2 (D) trained for clinical bactericidal outcomes. Combinations are plotted and shaped as in Figure 4. Regions are colored as in (B) based on decision tree classification thresholds learned during training.

There were too few drug combinations with clinical outcome scores to evaluate models with a held-out test set. Therefore, we trained random forest classifiers using the same approach applied for SOC and BPaL RMM and assessed their performance in cross-validation. We observed high performance (AUC > 0.8, similar to performances in the RMM) in models using many subsets of conditions (Table S7). Dormancy alone was a predictive condition (AUC=0.77, Table S7), suggesting that treating non-replicating Mtb is important for identifying effective drug combinations in humans. As with the BPaL predictions in the RMM, synergy in dormancy is associated with improved clinical outcomes (Table S8, log_2_FIC_50_ dormancy minimum, p=0.03, FDR p=0.41). The three-condition subset of butyrate+dormancy+cholesterol-high was the highest performing three-condition model (Table S7), suggesting that the information in these three conditions may be highly informative for understanding *in vivo* drug treatment that is not species-specific. Together, these results support using the *in vitro* data to train models that can be used to predict treatment outcomes in humans.

We generated predictions using the high-performing butyrate+dormancy+cholesterol-high model. We observed that candidate combinations had different predicted classifications for the SOC RMM and clinical models (Figure 5B), mirroring the discordance we previously noted (Figure 5A). Many of the candidate combinations using the twelve drugs were predicted to be +C1 for the clinical outcome (447, ∼79%, Figure 5B, Table S6), suggesting there may be many treatment improving combinations remaining to be tested using existing TB drugs. We compared the BPaL (RMM) and SOC (clinical) RF classifiers and found weak but significant correlation between their predictions (Figure S8B; Spearman’s rho=0.31, p<0.001). Notably, there were no combinations predicted (or annotated) to be better than BPaL in RMM that were also worse than SOC in the clinic, supporting the use of the RMM for identifying treatment improving combinations using BPaL as a benchmark.

To define a set of rules for rationally designing clinically effective drug combinations, we used the decision tree approach and generated two example rulesets (Figure 5C and D). The clinical rulesets require antagonism in butyrate and potency in a lipid-rich environment (or synergy in dormancy) (Figure 5C and D). Though it is not intuitive to choose combinations with antagonistic pairs, we previously identified high-order drug combination antagonism as important for classifying +C1 drug combinations (Larkins-Ford et al., 2021), supporting the notion that in some growth conditions, average *in vitro* drug pair antagonism may be associated with better outcomes relative to the SOC in both clinical and mouse studies. In other conditions, particularly in dormancy (Figure 3C, Figure 5D), synergy should be prioritized. For example, we observed synergy between isoniazid and pyrazinamide in dormancy (Table S1). This pair was recently observed to be synergistic in patients by PET-CT imaging in the first few weeks of clinical treatment (Xie et al., 2021)(Figure S7). For the SOC (clinical) DT, we note that HPZM, the intensive phase drug combination from Study 31 that shortens treatment time over the SOC in clinical trials (Dorman et al., 2021), satisfies one rule (antagonism in butyrate) but barely misses the threshold for the second rule (potency in cholesterol-high or synergy in dormancy), incorrectly predicting it to be -C1. Using RF and DT models, we predict that there are treatment shortening combinations among the drugs in this 12-drug set (Table S3, Figure 5C and D). We anticipate that as more clinical data become available, we will be able to refine the rule sets and improve prediction accuracy. Using *in vitro* drug combination measurements, rules for rationally designing drug combinations can be written that are interpretable and allow for comparisons between human and preclinical studies.

## Discussion

Our original goal in this study was to make the prediction of *in vivo* outcomes more efficient by factoring high-order combinations into more scalable parts: drug pairs. In the process, we found that models using pairwise *in vitro* measurements were predictive of high-order combination treatment in mouse models of disease relapse (RMM). Classifications based on pairwise instead of high-order measurements enabled us to predict three- and four-way drug combination outcomes *in vivo* among 12 TB antibiotics using around ten-fold fewer combination measurements than would have been required if we used direct DiaMOND measurement of each high-order combination. We increased the prediction resolution by introducing a higher threshold for classifications of *in vivo* outcomes (better than BPaL). With this approach, we predicted outcomes in 536 candidate combinations with no published RMM outcomes. We narrowed the number of combinations predicted to be top performers to 73 (14%) as better than BPaL in the RMM.

Factorization of high-order combinations into pairwise drug units also enabled us to develop predictive models that were very interpretable. We learned that a drug pair could be a building block upon which to assemble high-order combinations. We defined sets of rules that guide the construction of treatment-improving high-order combinations from effective building blocks. For example, one such rule is that one drug pair must be synergistic in the *in vitro* dormancy condition, and another pair must be potent in a growth medium where high levels of cholesterol are the carbon source. Though our machine learning models are more accurate than the rulesets from decision trees, these simple rules enabled us to rationally design combinations by evaluating systematic pairwise data. We applied the same classification and rule-building framework to outcomes in clinical trials, finding that the principles of combination design in the RMM translate to clinical outcomes. The clinical and RMM rulesets included pairs that are synergistic in dormancy and potent in lipid-rich conditions, consistent with notion that treating the most refractory bacteria in an infection is important for improving treatment outcomes. This is both intuitive and consistent with the decades of preclinical and clinical studies (Mitchison, 1996, Fox et al., 1999, Gengenbacher and Kaufmann, 2012, Kerantzas and Jacobs, 2017).

Constructing combinations from building block pairs is harmonious with the approaches used to design preclinical studies and clinical trials in that drugs are added to an effective base combination and evaluated as to whether or not the new combination improves outcomes (Fox et al., 1999, Gillespie et al., 2014, Bonnett et al., 2017, Rustomjee et al., 2008, Lee et al., 2019, Nuermberger et al., 2004a, Nuermberger et al., 2006, Nuermberger et al., 2008, Tasneen et al., 2015, Tasneen et al., 2011, Rosenthal et al., 2012, Tasneen et al., 2016). Using the available data, we predict that there are other treatment improving combinations in the existing drug combination space. In the future, the design principles will allow us to construct candidate combinations “at-a-glance” using practical and cost-effect pairwise combination measurement.

The rulesets we define establish a framework for combination design in experimentally tractable sizes: properties of a drug pair. We anticipated averaged pairwise data to predict a combination outcome. Instead, we found that properties defining the “best” (e.g., most potent or most interacting) pair in a combination were most informative. Ideally, each objective in a rule set should be achieved with a different pair. In this way, each pair can be viewed as a building block that not only enables us to construct combinations rationally from the bottom-up but also to identify how established combinations may be improved. Our initial rule sets assemble two pairs into 3-way combinations but generally leave a degree of freedom for choosing the fourth drug for 4-way combinations. We expect to define the third rule and enable 4-drug combination design when more 4-way *in vivo* data are available for model training.

The features used in each rule are also specific to the metric type, e.g., potency or interaction, allowing us to evaluate whether synergy is a requirement of the best combinations. One rule set for designing combinations that are better than BPaL in the RMM starts by assembling pairs with a synergistic pair in the dormancy *in vitro* model. Synergy separates combinations at the BPaL threshold, especially in dormancy. Classification around the standard of care is not driven by pairwise (this study) or high-order (Larkins-Ford et al., 2021) drug interactions. Furthermore, antagonism (not synergy) in *in vitro* models such as butyrate increases the likelihood of a combination performing better than the standard of care *in vivo*. These seemingly disparate rules may reflect the difference between simply achieving treatment efficacy (SOC) and improving treatment (better than BPaL) or may be indicative of which populations are easiest to sterilize (actively growing cells can be killed rapidly with drug pairs that are potent together) versus those where synergy is necessary (dormancy). To design a top-performing combination, having drugs that enhance the effect of each other could aid in targeting the most refractory cells in an infection (e.g., dormant/non-replicating) (Mitchison, 1996, Fox et al., 1999, Gengenbacher and Kaufmann, 2012, Kerantzas and Jacobs, 2017, Gold and Nathan, 2017, Saito et al., 2021). Further study is required to evaluate where these rulesets can be understood in the dynamic course of drug response in TB granulomas where Mtb residing in multiple compartments can respond differentially to treatment.

Drug sensitivity is highly dependent on the growth environment (Tarshis and Weed, 1953, Pethe et al., 2010, Zhang et al., 2013, Sanders et al., 2018, Sinclair et al., 2019, Cokol et al., 2019, Lamont et al., 2020). Therefore, the profiles of pairwise potencies and drug interactions were different in each of the seven *in vitro* growth models we evaluated. We constructed classifiers using data from each *in vitro* model alone and from all possible combinations of conditions to assess the relative importance of pairwise data from each growth environment. There was a strong predictive signal in most of the *in vitro* growth models. Still, the strongest signals were from the lipid-rich and dormancy-inducing conditions (butyrate, dormancy, cholesterol-high) when used together in classifiers. Some conditions provided orthogonal information that complemented and refined classifiers based on the top individual conditions (butyrate, dormancy, cholesterol-high), consistent with our understanding of the microenvironments where persisters reside, e.g., in lipid-rich environments and in conditions that lead to non-replicating subpopulations of Mtb (Fox et al., 1999, Daniel et al., 2011, Sarathy and Dartois, 2020, Gold and Nathan, 2017, Saito et al., 2021).

Predictive models are only as good as the data on which they are based, so our ability to make predictions and interpret combination design principles are dependent on the available *in vivo* and clinical combination outcome data. We anticipate that our classifiers will be refined and improved with future iterations that incorporate *in vivo* tests of combinations from our predictions and those that include antibiotics with new target profiles (Aldridge et al., 2021). The principles of ruleset design for the relapsing mouse model were generalizable to the clinical outcomes. As both classifiers are refined with accumulating data, we can compare rule sets to better define situations in which the RMM is predictive of clinical outcomes. These approaches will allow us to utilize best the rich information provided by preclinical and clinical studies through parallel *in vitro* studies, making bottom-up and top-down coordinated methods for the rational design of combination therapies for TB.

## Acknowledgments

We thank members of the Aldridge laboratory, K. Mdluli, and J. Silverman for their insightful discussion. This work was supported, in part, by the Bill & Melinda Gates Foundation OPP1189457. This work is supported in part by NIH grant 1U54CA225088: Systems Pharmacology of Therapeutic and Adverse Responses to Immune Checkpoint and Small Molecule Drugs for AS.

## Author contributions

JLF, NV, YD, and BBA conceived and designed the experiments. JLF, NV, and YD performed the experiments. JLF, NV, YD, AS, and BBA conceived and designed the computational analysis. JLF, NV, YD, and performed the computational analysis. The manuscript was written by JLF, YD, AS, and BBA. All authors contributed to the interpretation of the results and editing of the manuscript.

## Declaration of Interests

The authors declare no competing interests.

## Methods

### Mtb culturing and in vitro pairwise drug response measurements

To expand the pairwise drug combination response dataset from 10-drugs (described in (Larkins-Ford et al., 2021)) to 12-drugs, we use DiaMOND to measure 2-way dose-response curves with sutezolid and SQ109 against each other and the 10-drug set. The pairwise data with sutezolid is new to this study. Some of the SQ109 pairwise measures were reported in (Egbelowo et al., 2021), and the remaining combination measures in other growth environments are new to this study. All experiments were performed using the same procedures as described in (Larkins-Ford et al., 2021). Briefly, drug response was measured using an autoluminescent reporter strain of *M. tuberculosis* Erdman (transformed with a single copy chromosomal integration of pMV306hsp+LuxG13; (Andreu et al., 2010)), and metrics were averages of at least biological duplicate experiments. DiaMOND requires single- and equipotent drug combination dose responses to determine the potency and drug interactions. A 1.5-fold, ten-dose resolution dose-response was used for all experiments. SQ109 and sutezolid (non-metabolite form) were provided by Sequella, Inc. Drugs were stored and dispensed in DMSO using a digital drug dispenser (HP D300e).

The base medium of the standard and acidic *in vitro* models consisted of 7H9 Middlebrook medium supplemented with 10% OADC (0.5g/L oleic acid, 50g/L albumin, 20g/L dextrose, and 0.04g/L catalase), 0.05% Tween-80, and 25µg/mL kanamycin (to maintain selection of reporter-carrying Mtb). The base medium of the other *in vitro* models was 7H9 (4.73g/L) supplemented with fatty acid-free BSA (0.5g/L), NaCl (100mM), tyloxapol (0.05%), and 25µg/mL kanamycin. All *in vitro* model media were buffered to pH7.0 with 100 mM MOPS except acidic (buffered to pH5.7 with 100 mM MES). Carbon sources were added to *in vitro* model media to final concentration as follows: acidic and standard (glycerol, 0.2%), butyrate and dormancy (sodium butyrate, 5mM), valerate (valeric acid, 0.1%), cholesterol (cholesterol, 0.05mM), and cholesterol-high (cholesterol, 0.2 mM).

Mtb were cultured at 37℃ with aeration unless noted. Cells were maintained in standard media and passaged while in the mid-log phase (OD_600_ < 0.7). Mtb were acclimated to *in vitro* model growth medium for 2-6 doubling times prior to treatment for the DiaMOND assays. Acclimated Mtb were seeded at OD_600_=0.05 at 50uL per well onto 384-well plates with antibiotics pre-dispensed. The simple dormancy model is based on the butyrate medium, supplemented with sodium nitrate (5mM), sealed, and cultured without aeration to lower oxygen levels. After 28 days, dormant Mtb are plated (20uL per well) on antibiotic-seeded wells, the plates sealed and incubated. After seven days, 80uL of standard medium was added to each well, and plates were incubated with aeration for recovery and growth inhibition measurements.

Growth inhibition was measured by OD_600_ (for all conditions except dormancy) or luminescence (dormancy) using a Synergy Neo2 Hybrid Multi-Mode Reader (BioTek). The constant and terminal times are as follows in days, respectively: standard (4.2), acidic (6, 12), butyrate (6, 10), valerate (9, 15), cholesterol-high (12, 24), cholesterol (7, 28), dormancy (2, 4 into recovery). Growth inhibition measurements were processed, and dose-response metrics were calculated using custom scripts written in MATLAB (MathWorks).

### Drug pair dataset and data structure

For modeling and analysis, we used a 12-drug 2-way drug response dataset comprised of data from a 10-drug combination dose-response DiaMOND dataset (Larkins-Ford et al., 2021), a DiaMOND study of drug interactions with SQ109 (Egbelowo et al., 2021), and new measurements (sutezolid combinations and select SQ109 combinations). We selected the drug pair data from the dataset and used the dose-response metrics (AUC_25_, E_inf_, GR_inf_; higher (positive) AUC_25_ and E_inf_ values are potent and lower (negative) GR_inf_ values are potent) and drug interactions (log_2_FIC_50_, log_2_FIC_90_; negative and positive values indicate synergy and antagonism, respectively) from the constant and terminal time points. These drug pair metrics were aggregated via the minimum, maximum, and mean summary statistics for each high-order drug combination. Drugs with the same mechanism of action were excluded from any drug pair and high-order drug analysis (i.e., linezolid+sutezolid or rifampicin+rifapentine are not in candidate combinations).

### in vivo annotation of drug combinations

Annotations of drug combinations for the SOC outcome (+C1/-C1) were taken from a previous study (Larkins-Ford et al., 2021). The same studies were used to annotate the BPaL outcome (+C2/-C2). Twelve combinations were evaluated in Phase 2b trials for bactericidal activity using either culture negativity or time to positive (TTP) culture microbiological outcomes after eight weeks of treatment (Table S3). To increase the number of combinations for training machine learning models and because of the high clinical efficacy of bedaquiline-containing combinations, we also included one Phase 2a study (Diacon et al., 2015), where three bedaquiline-containing combinations (B+C+Pa+Z, B+C+Pa, B+C+Z) were tested for early bactericidal activity after 14 days of drug treatment using the TTP outcome. Similar conclusions can be drawn from comparing culture negativity (at eight weeks) and TTP (up to 56 days) (Phillips et al., 2016, Tweed et al., 2019, Olaru et al., 2014, de Knegt et al., 2017). We confirmed that including these combinations did not skew our candidate prediction results by comparing predictions to those made by a model that excluded these three combinations (R = 0.96 Pearson correlation, Figure S9).

### Data processing, computational analyses, and visualization

All data processing, computational analyses, and visualizations were performed in R (v4.0.1) using the tidyverse environment packages (v1.3.0). The readxls (v1.3.1) and openxlsx (v4.1.4) packages were used for data table import and export. The prcomp function from the stats package was used for PCA. Features with more than 35% missing data points were excluded from machine learning and PCA. Mean value imputation (Dray and Josse, 2014) was used for the remaining features with missing data. All features were mean-centered and scaled to unit variance prior to PCA. The ggplot2 (v3.3.0), ggpubr (v0.3.0), and ggrepel (v0.9.1) packages were used for all visualizations.

### Machine learning

All machine learning tasks, including model training and evaluation in cross-validation, were performed using the “machine learning in R” (mlr v2.17.0) package with individual learners loaded from additional packages (random forest, randomForestSRC (v2.9.3); Bayesian additive regression tree, bartMachine (v1.2.6); extreme gradient boosting, xgboost, (v1.4.1.1); k-nearest neighbor, kknn (v1.3.1); logistic regression, stats (v4.0.1); naïve Bayes, naiveBayes (v0.9.7); neural net, neuralnet (v1.44.2)). Models were evaluated on a 30% proportion of data (test) withheld from training. The test/training split was selected by random 30/70% partitioning of the data ten times and identifying a representative partition that had closest estimated model performance to the mean of the ten iterations (Table S10). Where appropriate, model performance was also estimated via cross-validation with a Monte-Carlo resampling strategy that partitioned the training (70% proportion) data into further 70/30% training/test splits across ten iterations. The Youden’s J (Youden, 1950) was used to select the optimal classification threshold based on training data.

### Decision tree and rule set determination

Decision trees were constructed in R using the rpart function (rpart package, v4.1-15), and rules and thresholds were analyzed using the rpart.plot package (v3.1.0). The minimum number of combinations for splitting a node was set to two, and the minimum terminal leaf size was set to five (RMM SOC and BPaL) or two (clinical SOC). Trees were allowed to grow fully.

### “Leave-one-drug-out” analysis

For each of the 12 drugs, annotated combinations containing that drug were withheld from model training. Models were trained with the remaining annotated combinations, and performance on data containing the withheld drug was determined.

### Statistical analysis

Statistical analyses were performed using the stats, ggpubr (v0.3.0), and rstatrix (v0.5.0) packages in R. Statistical significance threshold was chosen to be less than 0.05, unless otherwise indicated. The Wilcoxon rank-sum test was used to compare mean values across outcome groups. The Benjamini-Hochberg method was used to control the false discovery rate (FDR) for multiple hypothesis testing (Benjamini and Hochberg, 1995). Pearson’s correlation was used to measure linear correlations.

### Drug pair enrichment analysis

To determine if +C2 combinations contained signature sets of drugs, we tested for over-representation in the +C2 candidate drug combinations using Fisher’s Exact Test. We performed tests for each drug, drug pair, and three-drug combination and controlled the false discovery rate (FDR, (Benjamini and Hochberg, 1995))

## Supplemental Information

**Figure S1.** Separation of annotated drug combinations by PCA. Projection of the pairwise *in vitro* combination data from all *in vitro* models onto PCs 1 and 2 (top) and PCs 1 and 3(bottom). (A) Points are colored by outcome in the RMM: (A) blue=+C1, better than standard of care; red=-C1, standard of care or worse; (B) green=+C2, better than BPaL; orange=-C2, BPaL or worse. Percent variance explained by each PC indicated in the axis title. Outer box and whisker plots show the distributions of combination classes along each PC.

**Figure S2.** Model performance across different numbers of *in vitro* conditions. Scatter plots of model training AUC for SOC and BPaL classifiers for models trained with data from indicated number of *in vitro* conditions (one to seven). Median performance of every model is shown with black dashed lines (SOC AUC=0.83, BPaL AUC=0.88).

**Figure S3.** Top three-condition model performance and predictions. Test performance for the six highest performing three-condition models during training: (A) butyrate+standard+valerate, (B) dormancy+cholesterol-high + standard, (C) butyrate+dormancy+standard, (D) acidic+dormancy+cholesterol-high, (E) acidic+cholesterol+dormancy, (F) butyrate + dormancy + cholesterol-high. ROC (top) and PR (bottom) curves are labeled as in Figure 2A. Probability scatter plots are on the right and labeled as in Figure 2D.

**Figure S4.** Contribution of conditions to model performance. (A) Scatter plots of training performance for models without the indicated condition compared to models including the indicated condition. Change in model performance by inclusion of the condition is indicated by color (increased (blue), decreased (red), or indifferent (grey)). Dashed line indicates the line of “indifference”, where model performance does not change with or without indicated condition. Single condition training performance indicated above plot and with solid line. Percentage of models with increased or decreased performance are shown. (B) Model performance density plot of models with (green) and without (red) butyrate+dormancy+cholesterol-high (red).

**Figure S5.** Univariate analysis of features using combined training and test data. Univariate feature analysis in SOC and BPaL models. (A) Box plots showing the distribution of values for drug interaction (log_2_FIC_50_ and log_2_FIC_90_), and drug potency (E_inf_, GR_inf_, and AUC_25_ based on BPaL (green = +C2 and yellow = -C2) outcome. (B) Scatter plot of p-values for the Wilcoxon Rank Sum test evaluated for predicting SOC (-C1 vs +C1) and BPaL (-C2 vs +C2) outcomes. Features are colored by *in vitro* condition and shaped by metric type (circle, AUC_25_; square, E_inf_; downward triangle, GR_inf_). P-values are corrected for multiple hypothesis testing within each outcome group (e.g., corrected for SOC comparison separate from BPaL comparison). Dashed lines show p=0.05. Features with FDR p-values <0.05 are annotated with extra information such as time (C or T for constant or terminal, respectively) and the summary statistic type (min, mean, or max). Linear regression line (solid black), confidence interval (shaded region), Pearson correlation coefficient (R) and associated p-value are indicated on plot. (C) Scatter plot of p-values from the Wilcoxon rank-sum tests contrasting values of individual drug interaction features across SOC (-C1 vs. +C1) and BPaL (-C2 vs. +C2) outcomes. Plot elements are analogous to those in panel B. Features are shaped by metric type (upward triangle, log_2_FIC_90_; diamond, log_2_FIC_50_).

**Figure S6.** Alternative ruleset scatter plots. Scatter plots of two metrics from each subset of conditions identified to be important for outperforming BPaL for each subset of conditions: (A) butyrate+dormancy+choleterol-high, (B and C) butyrate+standard+valerate, (D and E) acidic+dormancy+cholesterol-high. Plots are labeled as in Figure 4. Combinations are separated into those that were used in decision tree model training (circle, top-left), testing (triangle, top-right), or are candidates (square, bottom-left). Selected drug combinations are indicated with labels. Plot regions are colored based on the decision tree classification using thresholds (dashed lines) learned during training. Selected drug pair metric values are indicated along plot margins. Logic formatted rules are written in the bottom-right of each panel.

**Figure S7.** “At-a-glance” drug pair *in vitro* metric heatmaps. Heatmap of drug pair data for selected drug interaction (A, C, E) and potency (B, D, F) features for the conditions butyrate (A, B), dormancy (C, D) and cholesterol-high (E, F). Drugs are indicated along plot margin using abbreviations as in Table 1. Drug pair data are colored by their values for the indicated metric and condition.

**Figure S8.** RMM predictions help stratify clinical SOC predictions. (A) Overlap in drug combination categorization between BPaL (RMM) and SOC (clinical) outcomes. Green/Blue and Orange/Red squares indicate treatment improvement agreement (+C2/+C1) between outcomes. Yellow/Blue and Green/Red squares highlight treatment improvement differences between outcome annotation (+C2/-C1 or -C2/+C1). (B) Probability scatter plot for BPaL model predictions (+C2 probability) and clinical model predictions (+C1 probability) using the butyrate+dormancy+cholesterol-high condition data. Annotated combinations are colored by clinical outcome when treatment improvement agrees, or split color is shown as in panel A. Model training combinations for both BPaL and clinical are labeled with circles. Combinations used for testing the BPaL model and training the clinical model training are labeled with diamonds. Candidate combinations (without annotations) are labeled with grey squares, and the number and percent of candidates in quadrants are indicated. Probability regions and threshold are labeled as in Figure 5 with regions colored for clinical outcome as in panel A

**Figure S9.** Prediction correlation from clinical models with and without Phase 2a trial combinations. Scatter plot of prediction probabilities from model trained with only Phase 2b trial combinations (12 combinations) and model trained with Phase 2a and Phase 2b trial combinations (15 combinations). Annotated combinations used for model training are indicated with circles (Phase 2b) and triangles (Phase 2a). Candidate combinations are in grey boxes. Linear regression line, Pearson correlation coefficient (R), and associated p-value are shown.

**Table S1.** Pairwise drug combination metric data.

**Table S2.** High-order drug combination summarized pairwise dataset

**Table S3.** Drug combination *in vivo* outcome annotations.

**Table S4.** ML algorithm performances.

**Table S5.** “Leave-one-drug-out” model performances.

**Table S6.** RF model predictions.

**Table S7.** Conditions subset model performances

**Table S8.** Univariate analysis of features for drug combination class separation

**Table S9.** Drug pair enrichment analysis.

**Table S10.** RF model performance with repeated training and test resampling.

## Notes

### Competing Interest Statement

The authors have declared no competing interest.

