## Supplemental Information for "Design principles to assemble drug combinations for effective tuberculosis therapy using interpretable pairwise drug response measurements"

### Figure S1

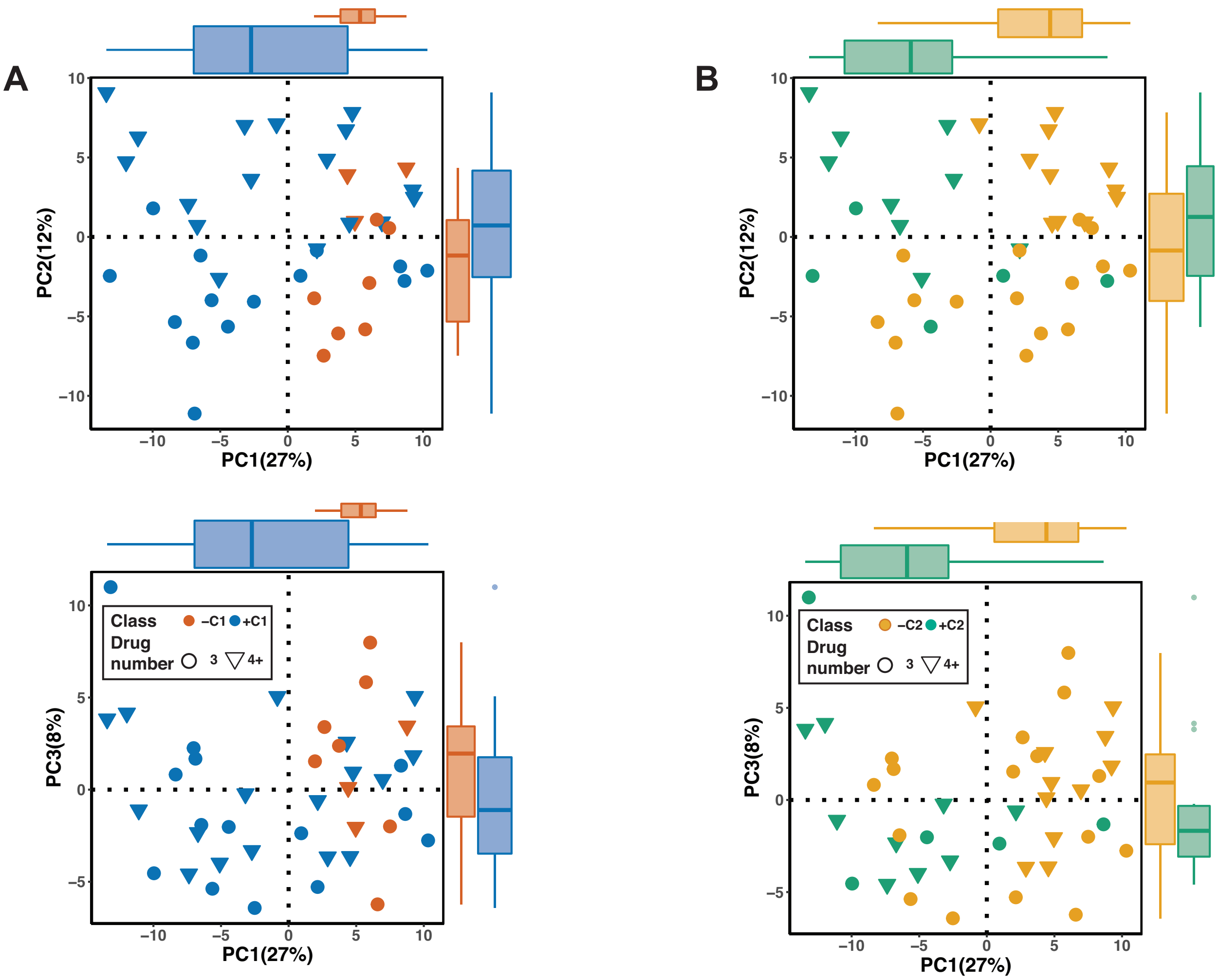


**Figure S1.** Separation of annotated drug combinations by PCA. Projection of the pairwise *in vitro* combination data from all *in vitro* models onto PCs 1 and 2 (top) and PCs 1 and 3(bottom). (A) Points are colored by outcome in the RMM: (A) blue=+C1, better than standard of care; red=-C1, standard of care or worse; (B) green=+C2, better than BPaL; orange=-C2, BPaL or worse. Percent variance explained by each PC indicated in the axis title. Outer box and whisker plots show the distributions of combination classes along each PC.

### Figure S2

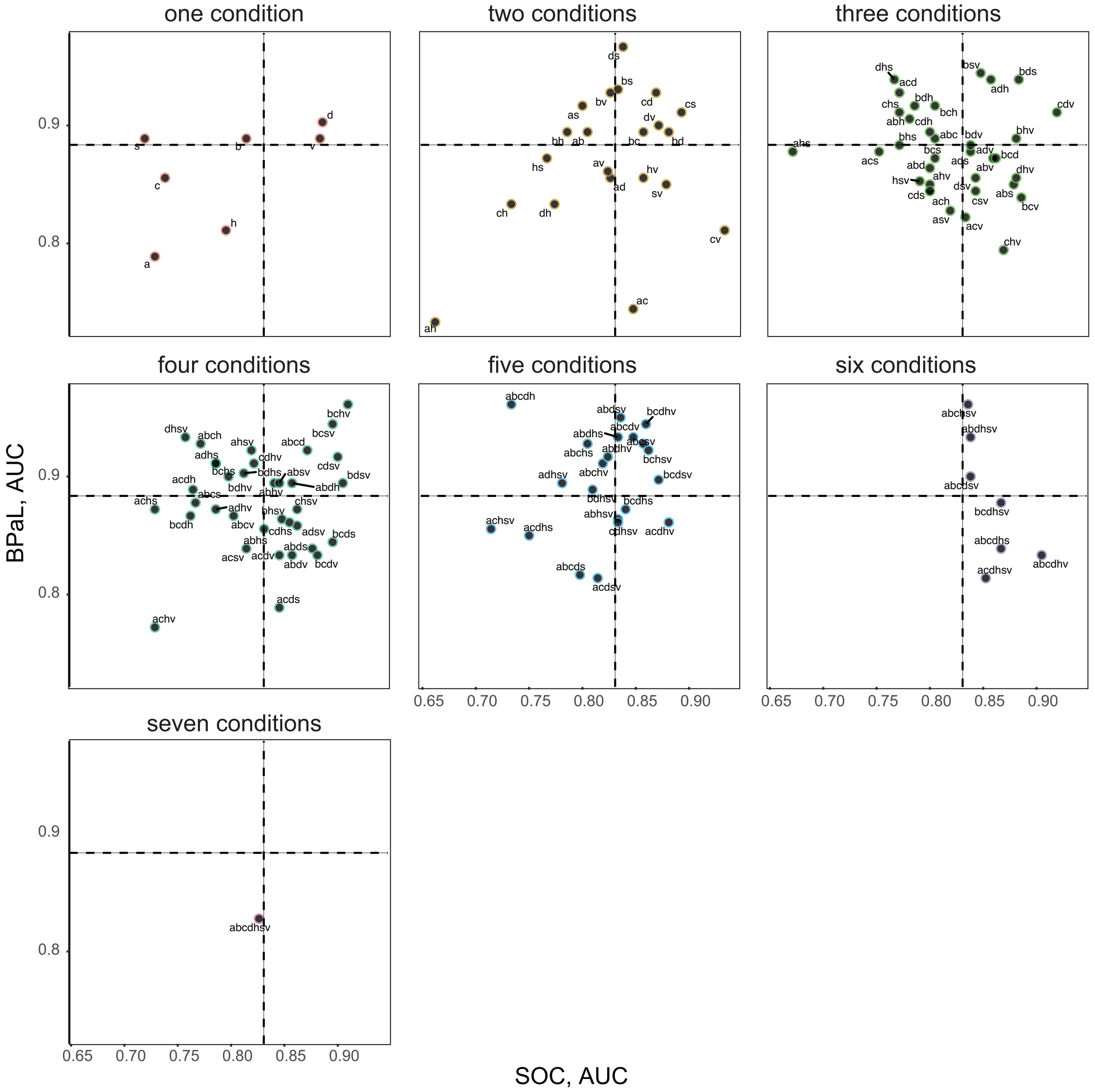


**Figure S2.** Model performance across different numbers of *in vitro* conditions. Scatter plots of model training AUC for SOC and BPaL classifiers for models trained with data from indicated number of *in vitro* conditions (one to seven). Median performance of every model is shown with black dashed lines (SOC AUC=0.83, BPaL AUC=0.88).

### Figure S3

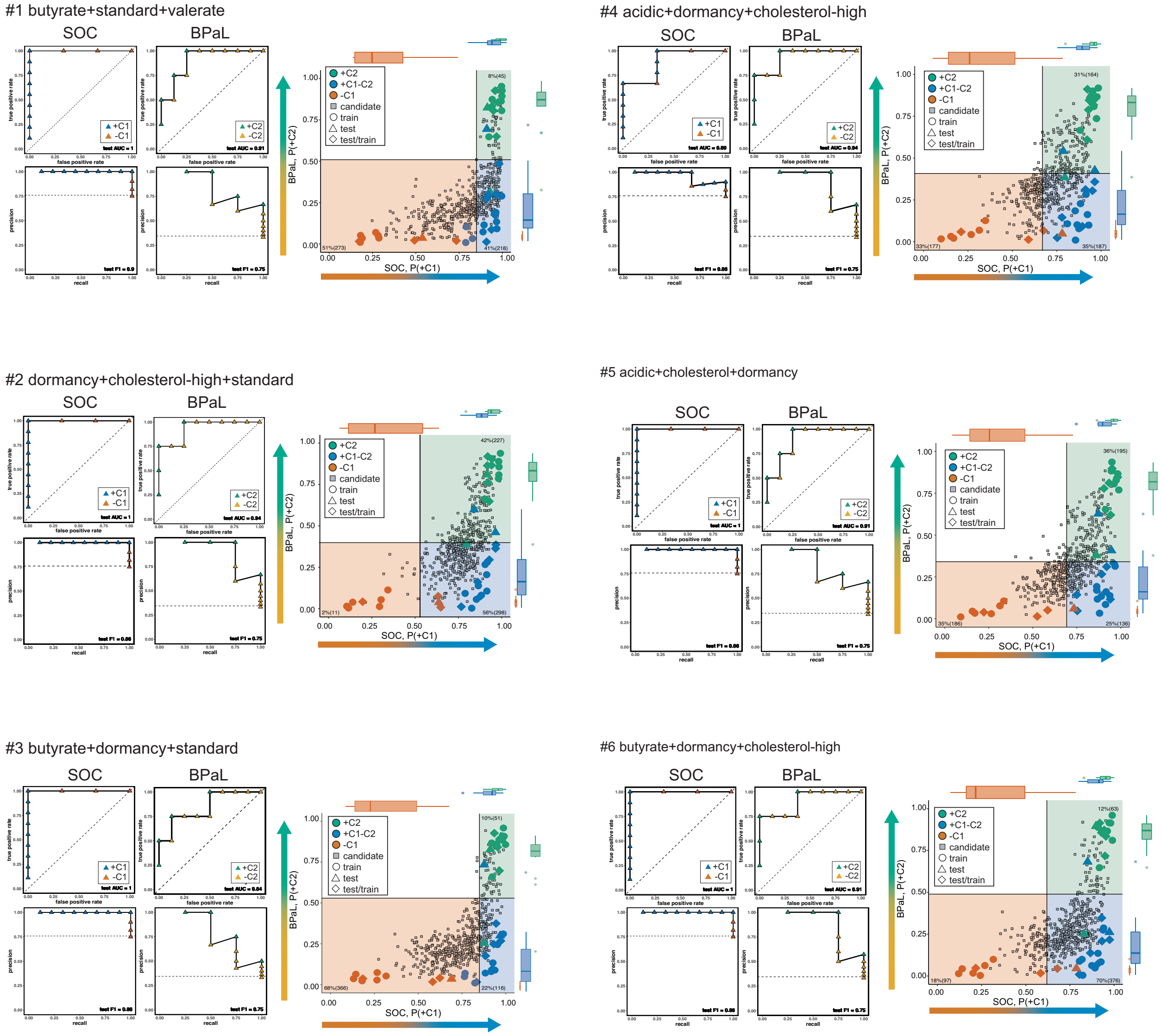


**Figure S3.** Top three-condition model performance and predictions. Test performance for the six highest performing three-condition models during training: (A) butyrate+standard+valerate, (B) dormancy+cholesterol-high + standard, (C) butyrate+dormancy+standard, (D) acidic+dormancy+cholesterol-high, (E) acidic+cholesterol+dormancy, (F) butyrate + dormancy + cholesterol-high. ROC (top) and PR (bottom) curves are labeled as in Figure 2A. Probability scatter plots are on the right and labeled as in Figure 2D.

### Figure S4

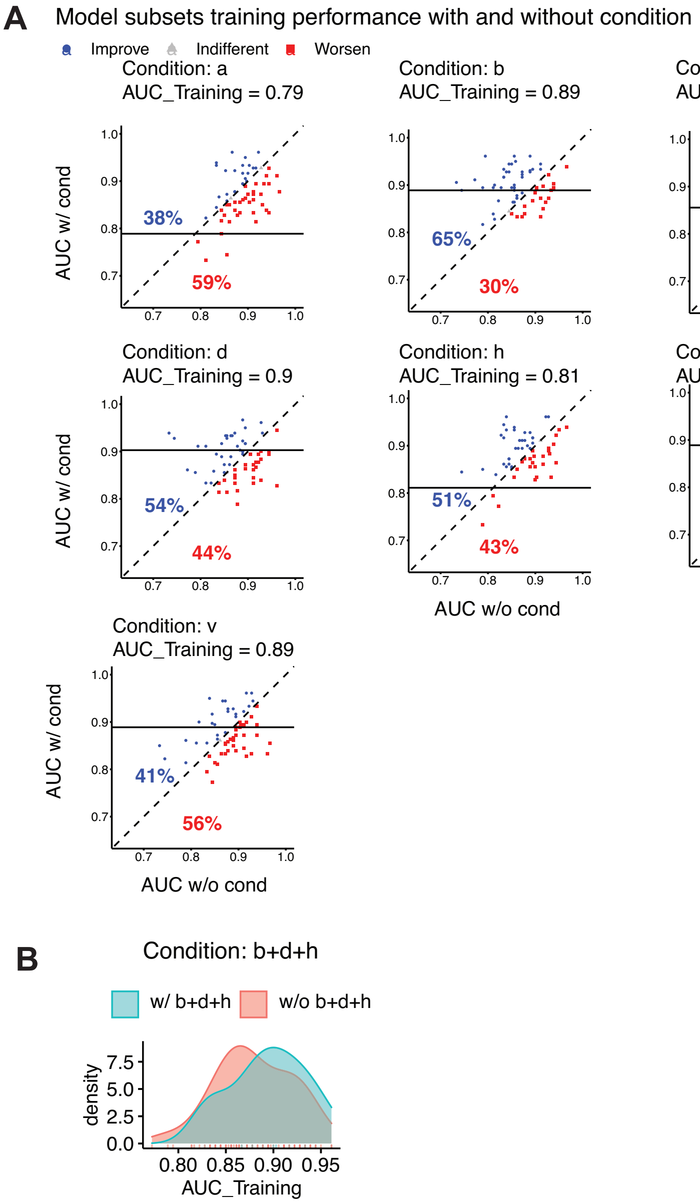


**Figure S4.** Contribution of conditions to model performance. (A) Scatter plots of training performance for models without the indicated condition compared to models including the indicated condition. Change in model performance by inclusion of the condition is indicated by color (increased (blue), decreased (red), or indifferent (grey)). Dashed line indicates the line of “indifference”, where model performance does not change with or without indicated condition. Single condition training performance indicated above plot and with solid line. Percentage of models with increased or decreased performance are shown. (B) Model performance density plot of models with (green) and without (red) butyrate+dormancy+cholesterol-high (red).

**Figure S5**

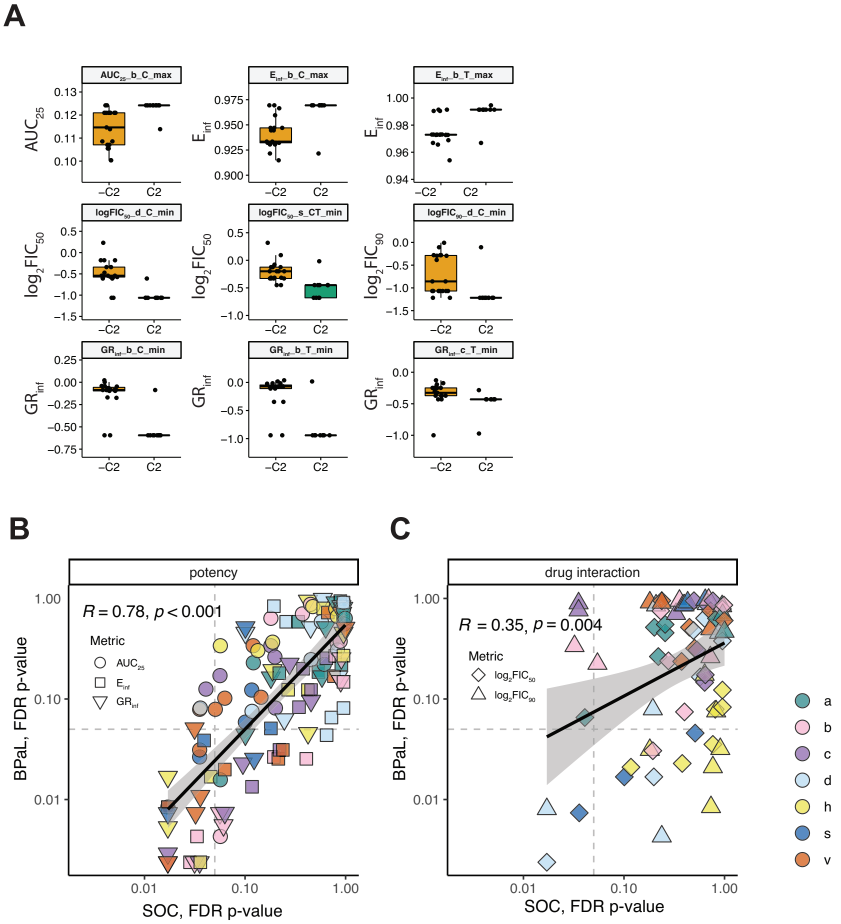


**Figure S5.** Univariate analysis of features using combined training and test data. Univariate feature analysis in SOC and BPaL models. (A) Box plots showing the distribution of values for drug interaction ($log_2FIC_{50}$ and $log_2FIC_{90}$), and drug potency ($E_{inf}$, $GR_{inf}$, and $AUC_{25}$ based on BPaL (green = +C2 and yellow = -C2) outcome. (B) Scatter plot of p-values for the Wilcoxon Rank Sum test evaluated for predicting SOC (-C1 vs +C1) and BPaL (-C2 vs +C2) outcomes. Features are colored by *in vitro* condition and shaped by metric type (circle, $AUC_{25}$; square, $E_{inf}$; downward triangle, $GR_{inf}$). P-values are corrected for multiple hypothesis testing within each outcome group (e.g., corrected for SOC comparison separate from BPaL comparison). Dashed lines show p=0.05. Features with FDR p-values <0.05 are annotated with extra information such as time (C or T for constant or terminal, respectively) and the summary statistic type (min, mean, or max). Linear regression line (solid black), confidence interval (shaded region), Pearson correlation coefficient (R) and associated p-value are indicated on plot. (C) Scatter plot of p-values from the Wilcoxon rank-sum tests contrasting values of individual drug interaction features across SOC (-C1 vs. +C1) and BPaL (-C2 vs. +C2) outcomes. Plot elements are analogous to those in panel B. Features are shaped by metric type (upward triangle, $log_2FIC_{90}$; diamond, $log_2FIC_{50}$).

### Figure S6

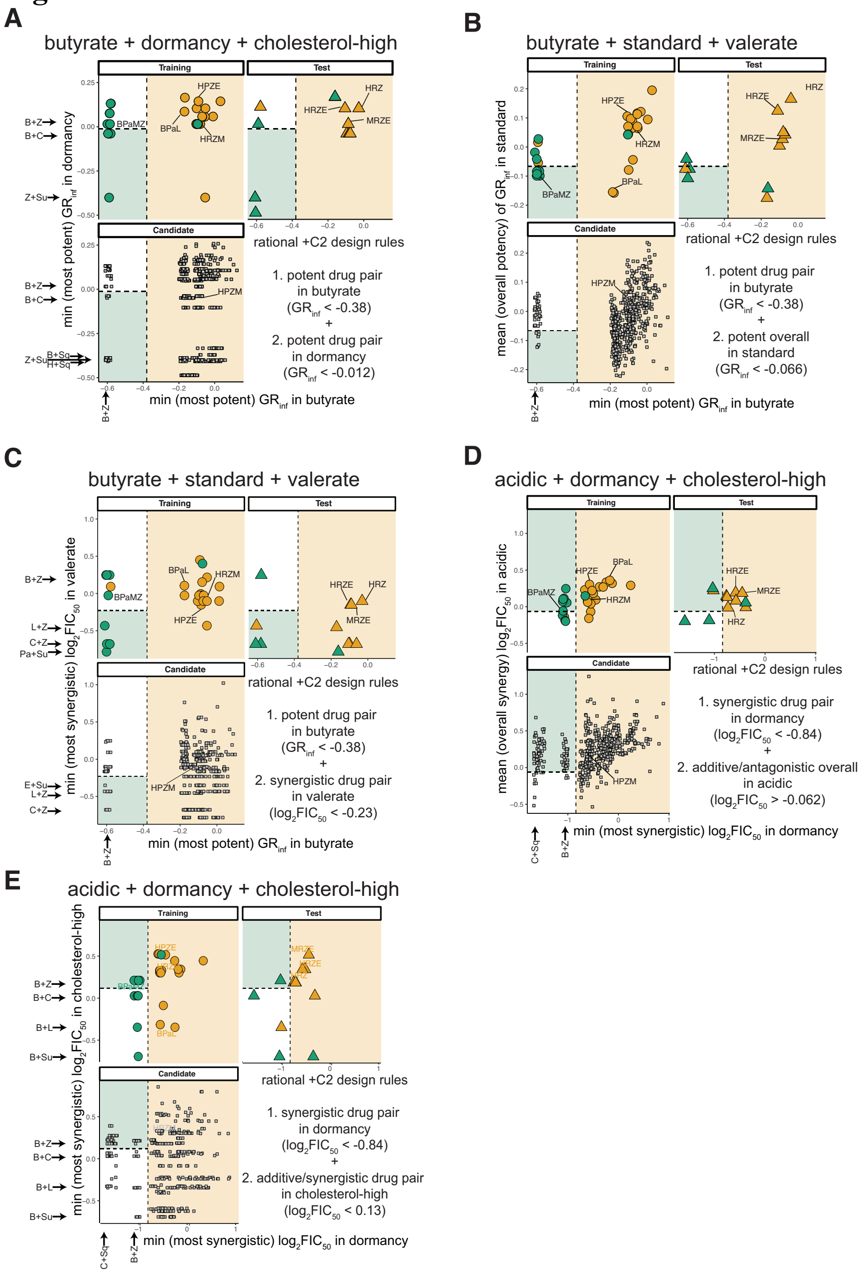


**Figure S6.** Alternative ruleset scatter plots. Scatter plots of two metrics from each subset of conditions identified to be important for outperforming BPaL for each subset of conditions: (A) butyrate+dormancy+choleterol-high, (B and C) butyrate+standard+valerate, (D and E) acidic+dormancy+cholesterol-high. Plots are labeled as in Figure 4. Combinations are separated into those that were used in decision tree model training (circle, top-left), testing (triangle, top-right), or are candidates (square, bottom-left). Selected drug combinations are indicated with labels. Plot regions are colored based on the decision tree classification using thresholds (dashed lines) learned during training. Selected drug pair metric values are indicated along plot margins. Logic formatted rules are written in the bottom-right of each panel.

### Figure S7

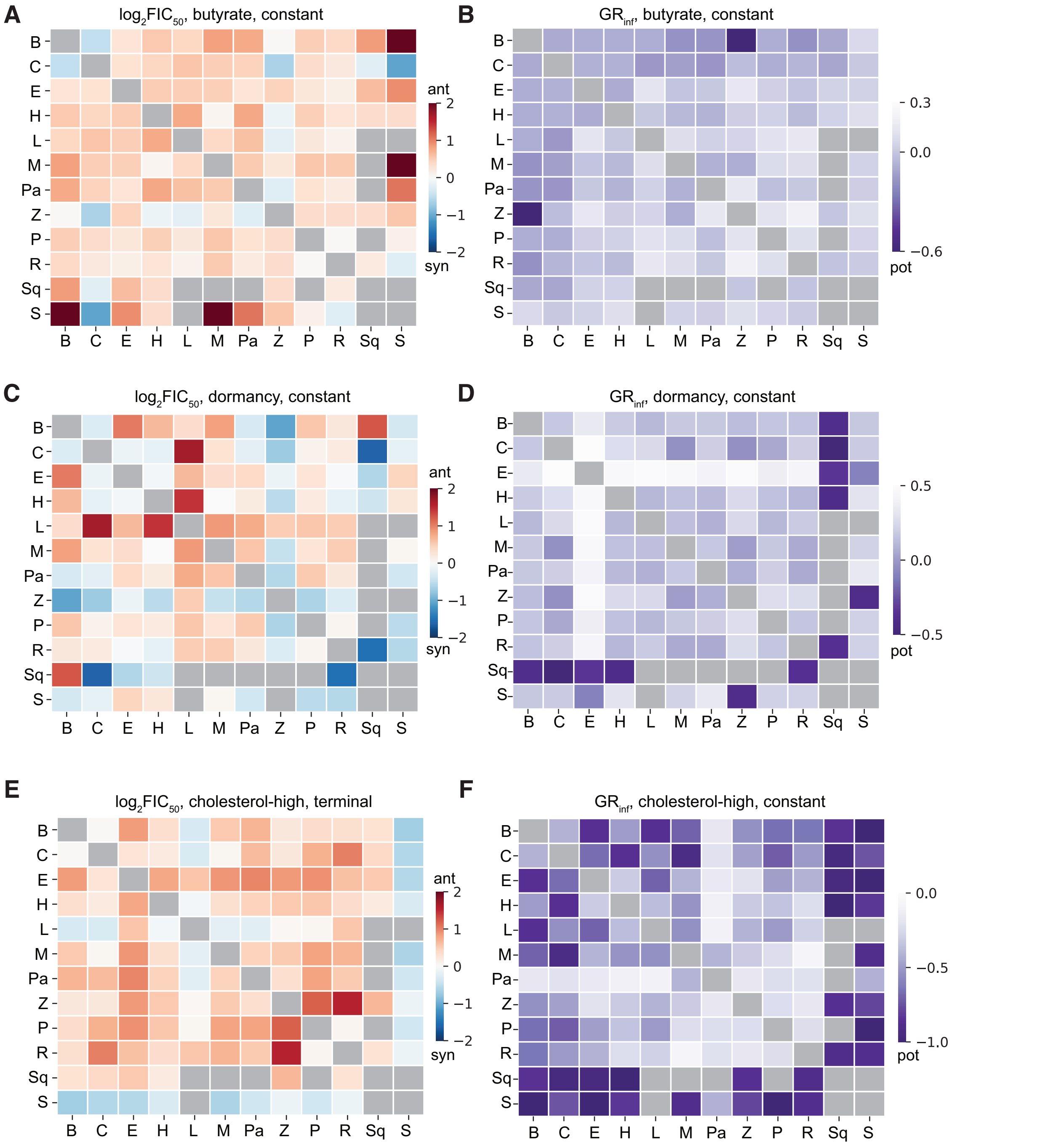


**Figure S7.** "At-a-glance" drug pair *in vitro* metric heatmaps. Heatmap of drug pair data for selected drug interaction (A, C, E) and potency (B, D, F) features for the conditions butyrate (A, B), dormancy (C, D) and cholesterol-high (E, F). Drugs are indicated along plot margin using abbreviations as in Table 1. Drug pair data are colored by their values for the indicated metric and condition.

**Figure S8**

A

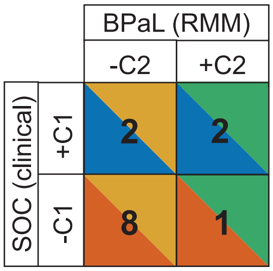


B

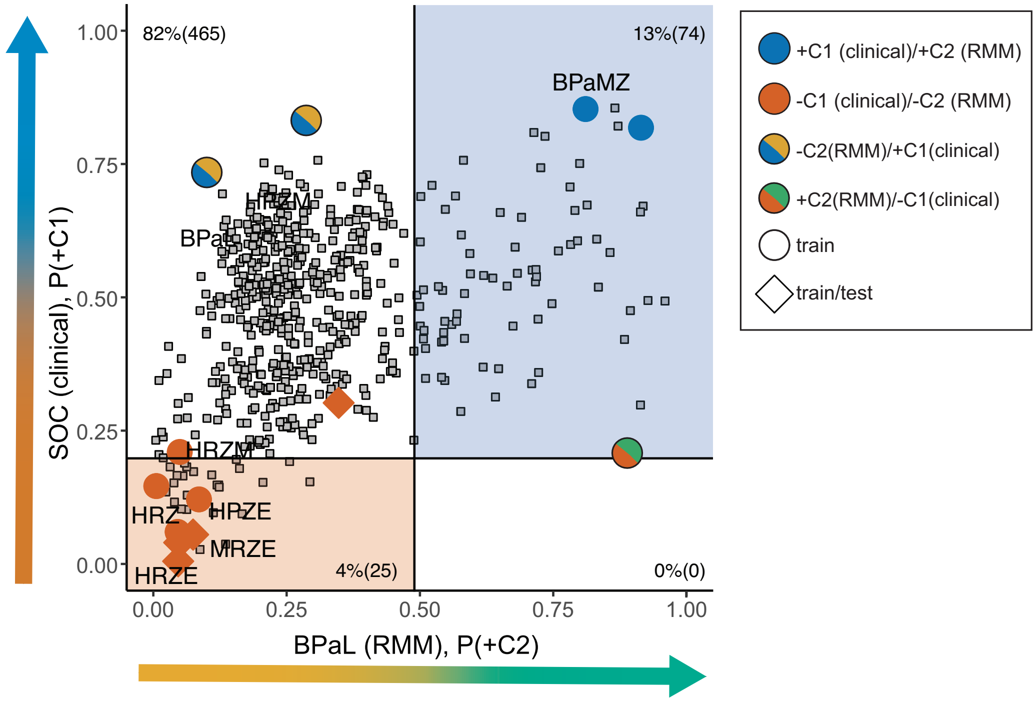


**Figure S8.** RMM predictions help stratify clinical SOC predictions. (A) Overlap in drug combination categorization between BPaL (RMM) and SOC (clinical) outcomes. Green/Blue and Orange/Red squares indicate treatment improvement agreement (+C2/+C1) between outcomes. Yellow/Blue and Green/Red squares highlight treatment improvement differences between outcome annotation (+C2/-C1 or -C2/+C1). (B) Probability scatter plot for BPaL model predictions (+C2 probability) and clinical model predictions (+C1 probability) using the butyrate+dormancy+cholesterol-high condition data. Annotated combinations are colored by clinical outcome when treatment improvement agrees, or split color is shown as in panel A. Model training combinations for both BPaL and clinical are labeled with circles. Combinations used for testing the BPaL model and training the clinical model training are labeled with diamonds. Candidate combinations (without annotations) are labeled with grey squares, and the number and percent of candidates in quadrants are indicated. Probability regions and threshold are labeled as in Figure 5 with regions colored for clinical outcome as in panel A.

**Figure S9**

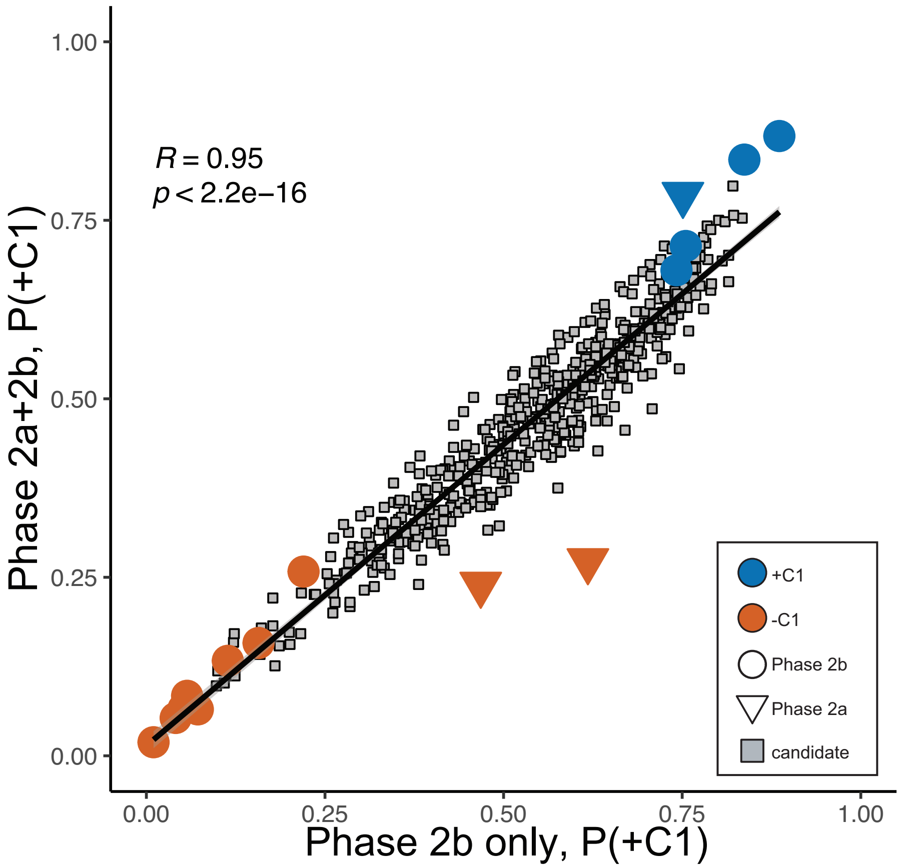


**Figure S9.** Prediction correlation from clinical models with and without Phase 2a trial combinations. Scatter plot of prediction probabilities from model trained with only Phase 2b trial combinations (12 combinations) and model trained with Phase 2a and Phase 2b trial combinations (15 combinations). Annotated combinations used for model training are indicated with circles (Phase 2b) and triangles (Phase 2a). Candidate combinations are in grey boxes. Linear regression line, Pearson correlation coefficient (R), and associated p-value are shown.
